# PLK1-Mediated Phosphorylation Cascade Activates the Mis18 Complex to Ensure Centromere Inheritance

**DOI:** 10.1101/2024.02.23.581399

**Authors:** Pragya Parashara, Bethan Medina-Pritchard, Maria Alba Abad, Paula Sotelo-Parrilla, Reshma Thamkachy, David Grundei, Juan Zou, Vimal Das, Zhaoyue Yan, David A. Kelly, Toni McHugh, Juri Rappsilber, A. Arockia Jeyaprakash

## Abstract

Accurate chromosome segregation requires the attachment of spindle microtubules to centromeres, which are epigenetically defined by the enrichment of CENP-A nucleosomes. During DNA replication, existing CENP-A nucleosomes undergo dilution as they get redistributed among the two DNA strands. To preserve centromere identity, CENP-A levels must be restored in a cell-cycle controlled manner orchestrated by the Mis18 complex. Here we provide a comprehensive mechanistic basis for PLK1-mediated licensing of CENP-A loading. We demonstrate that PLK1 interacts with Mis18α and Mis18BP1 subunits of the Mis18 complex by recognising self-primed phosphorylations of Mis18α (S54) and Mis18BP1 (T78 and S93) through its Polo-box binding domain. Disrupting these PLK1 phosphorylations perturbed the centromere recruitment of HJURP and new CENP-A loading. Biochemical and functional analyses show that phosphorylation of Mis18α and subsequent PLK1 binding is required to activate the Mis18α/β complex for robust Mis18α/β-HJURP interaction. Thus, our study reveals key molecular events underpinning the licensing role of PLK1 in ensuring accurate centromere inheritance.

**One-Sentence Summary:** PLK1 phosphorylation cascade licenses CENP-A loading by facilitating HJURP centromere recruitment via Mis18α/β activation.

## Introduction

The centromere is a key chromosomal locus that acts as a microtubule attachment site essential for the faithful segregation of genetic material to the daughter cells during cell division. In most eukaryotes, the centromere is epigenetically defined by a ∼50-fold enrichment of nucleosomes containing CENP-A, a histone H3 variant, compared to the rest of the genome (*1, 2*). During DNA replication, CENP-A nucleosomes are distributed between the old and newly replicated DNA, reducing the levels of centromeric CENP-A by half. To preserve centromere identity, a precise amount of CENP-A must be accurately reloaded onto the centromere at the correct time (*3, 4*). The loss of centromere identity or defective centromere formation results in chromosome missegregation or fragmentation, leading to aneuploidy and chromosome instability (*2, 5, 6*). In humans, the replenishment of CENP-A is enabled by the Mis18 complex (comprising of the Mis18α/Mis18β complex and Mis18BP1), which associates with the centromere during late mitosis/early G1 via interactions with the components of the Constitutive Centromere Associated Network (CCAN), including CENP-C and CENP-I (*7-12*). The centromere-associated Mis18 complex recruits the CENP-A specific chaperone HJURP, bound to CENP-A/Histone H4, to deposit CENP-A in G1 (*13-15*). The temporal restriction of CENP-A deposition to G1 is primarily regulated by the Cyclin Dependent Kinase 1 and 2 (CDKs) and PLK1 (*16-19*).

The Mis18α/β complex forms a hetero-hexamer of 4 Mis18α and 2 Mis18β. Two copies of Mis18BP1, via their N-terminal 130 amino acids, associate with the Mis18α/β hexamer to form a hetero-octameric complex (*16, 17*). CDKs control the timing of Mis18 complex assembly by phosphorylating specific residues on Mis18BP1 (T40 and S110), which inhibits Mis18BP1’s binding to the Mis18α/β complex. This prevents premature CENP-A loading until the end of mitosis (*16, 17*). CDKs also phosphorylate Mis18BP1 (T653) and HJURP (S210, S211 and S412) to disrupt their centromere localisation (*20*). While CDKs act as negative regulators of CENP-A deposition by disrupting the assembly of the Mis18 complex and its centromere association, PLK1 is suggested to play a positive regulatory role by promoting the centromere association of the Mis18 complex. PLK1 localises to the centromere at G1 in a Mis18 complex-dependent manner (*19*) and is proposed to license CENP-A deposition by facilitating the centromere association of the Mis18 complex through Mis18BP1 phosphorylation (*19*).

However, how PLK1 interacts with the Mis18 complex and what the roles of PLK1 phosphorylation of the Mis18 complex are in facilitating CENP-A loading have remained key outstanding questions for nearly a decade. Here, we show that PLK1 associates with the Mis18 complex by directly interacting with self-primed phosphorylation sites on Mis18α and Mis18BP1. Our biochemical, structural, and cellular functional studies reveal that a PLK1-mediated phosphorylation cascade regulates HJURP centromere recruitment and new CENP-A loading by regulating the Mis18α/β interaction with HJURP through conformational activation of the Mis18α/β complex.

## Results

### PLK1 directly interacts with Mis18α/β and Mis18BP1 in a phosphorylation dependent manner

McKinley and Cheeseman previously reported that PLK1 phosphorylation of the Mis18 complex, primarily Mis18BP1, is crucial for CENP-A deposition (19). To investigate the molecular basis for Mis18 complex-PLK1 interaction, we first probed whether PLK1 could directly interact with Mis18α/β and Mis18BP1 in vitro. Size exclusion chromatography (SEC) analysis indicated a weak interaction between Mis18α/β and PLK1 (Fig. 1A & 1B, black profile). Remarkably, when the Mis18α/β-PLK1 mix was incubated with ATP/MgCl_2_ to allow phosphorylation of Mis18α/β by PLK1, a robust Mis18α/β-PLK1 complex was formed (Fig. 1B, red profile). Similarly, His-MBP-Mis18BP1_1-490_ and PLK1 formed a robust complex only after PLK1 phosphorylation of His-MBP-Mis18BP1_1-490_ (Fig. 1C). We then assessed if the Mis18 complex consisting of both Mis18α/β and Mis18BP1_1-490_ could interact with PLK1 and we observed complex formation only upon the PLK1 phosphorylation of the Mis18 complex (Fig. S1A, black and red profiles). PLK1 employs its Polo-Box Domain (PBD) (Fig. 1A) to recognise substrates primed either by CDKs or by itself (*21, 22*). Our SEC and amylose pull-down analyses confirmed that PLK1 interacts with Mis18α/β, Mis18BP1_1-490_ and the Mis18 complex by recognising self-primed phosphorylations via PLK1_PBD_ (Fig. S1B-D).

**Fig. 1.**
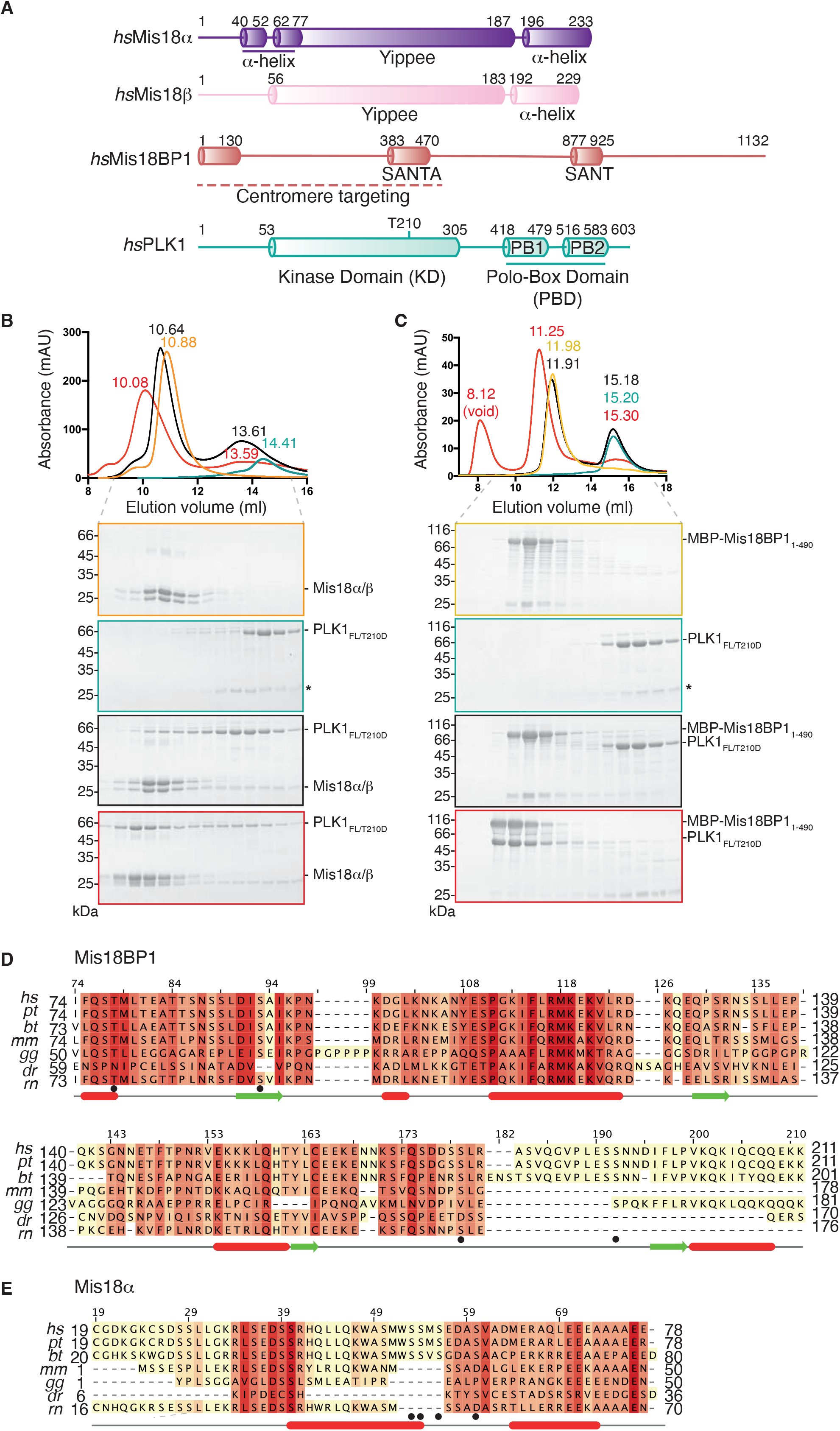
Mis18α/β and Mis18BP1 Interacts with PKL1 in a Phospho-Dependent Manner. (**A**) Domain architecture of Mis18α (purple) Mis18β (pink) Mis18BP1 (salmon) and PLK1 (green). (**B**) SEC profiles and corresponding SDS-PAGE analysis of Mis18α/β (orange) and PLK1 (green) individually, mixed together (black) and mix together with ATP/Mg^2+^ and incubated to allow phosphorylation (red). Asterisk denotes contaminant from the PLK1 purification. (**C**) SEC profiles and corresponding SDS-PAGE analysis of His-MBP-Mis18BP1_1-490_ (gold) and PLK1 (green) individually, mixed together (black) and mix together with ATP/Mg^2+^ and incubated to allow phosphorylation (red). Asterisk denotes contaminant from the PLK1 purification. (**D-E**) Multiple sequence alignment for and (**D**) Mis18BP1 and (**E**) Mis18α using MUSCLE (*46*) visualised with Jalview (*47*) with sequences from *Homo sapiens* (*hs*), *Pan troglodytes* (*pt*), *Bos taurus* (*bt*), *Mus musculus* (*mm*), *Gallus gallus* (*gg*), *Danio rerio* (*dr*) and *Rattus norvegicus* (*rn*). Secondary structure prediction was performed using JPred Secondary Structure Prediction (*48*). Black dots indicate phosphorylated residues identified by mass spectrometry.

Next, we aimed to identify the specific amino acid residues of the Mis18 complex phosphorylated by PLK1. Through mass spectrometry (MS) analysis on recombinantly purified Mis18α/β, Mis18BP1 and Mis18α/β/Mis18BP1 samples phosphorylated by PLK1, we identified phosphorylations that were then filtered by the presence/absence of the PLK1_PBD_ binding motif (S-S/Tph) and evolutionary conservation. In line with McKinley and Cheeseman (*19*), we identified phosphorylations on Mis18BP1 amino acid residues S93, S179 and S192 (Fig. 1D & Supplementary Table 1). Additionally, in the Mis18α/β/Mis18BP1 sample, we discovered previously unreported phosphorylation on Mis18BP1 T78. Both T78 and S93 are located in the N-terminal Mis18α/β-binding region of Mis18BP1, while S178 and S192 are within the unstructured region between the N-terminal Mis18α/β-binding and the SANTA domains of Mis18BP1 (Fig. 1D, black dots & Supplementary Table 1). Our MS data also revealed four amino acid residues in the N-terminus of Mis18α: S53, S54, S56 and S60 that were phosphorylated by PLK1 (Fig. 1E, black dots & Supplementary Table 1; also confirmed by (*19*)). Notably, these residues are positioned in the N-terminal region of Mis18α, which we have shown recently to fold back and interact with the Mis18α/β C-terminal α-helices (*23*) implicated in HJURP binding (*24, 25*). This suggests that PLK1 recognises Mis18α/β and Mis18BP1 through self-priming phosphorylation of residues located in regions involved in crucial protein-protein interactions.

### Structural basis for PLK1 recruitment to the centromere via the Mis18 complex

After confirming that phosphorylation of Mis18α/β and Mis18BP1 is essential for robust interaction with PLK1, we focused on identifying which phosphorylated residues are critical for this interaction. Mutating S53, S54, S56 and S60 of Mis18α to non-phosphorylatable alanine (Mis18α_4A_) abolished PLK1 binding (Fig. S2A). Further analysis with single point mutations helped narrow down the key PLK1-interacting residue in Mis18α to S54 (Mis18α_S54A_) (Fig. 2A, red profile). The SDS-PAGE migration pattern showed that the Mis18α_S54A_ mutant was still phosphorylated by PLK1, confirming that additional Mis18α residues undergo phosphorylation but are not essential for Mis18α-PLK1 interaction (Fig. 2A). In the case of Mis18BP1, making T78 and S93 non-phosphorylatable (Mis18BP1_T78A/S93A_) abolished PLK1 interaction, as shown by amylose pull-down assays and SEC analysis (Fig. S2B & Fig. 2B, red profile), and also significantly reduced PLK1 phosphorylation of Mis18BP1. Combining these mutations (Mis18α_S54A_ together with Mis18BP1_T78/S93A_) in the context of the Mis18 complex did not affect Mis18 complex formation but are needed to completely abolished PLK1 interaction (Fig. S2C, orange and red panels, respectively. Fig. S2D-E).

**Fig. 2.**
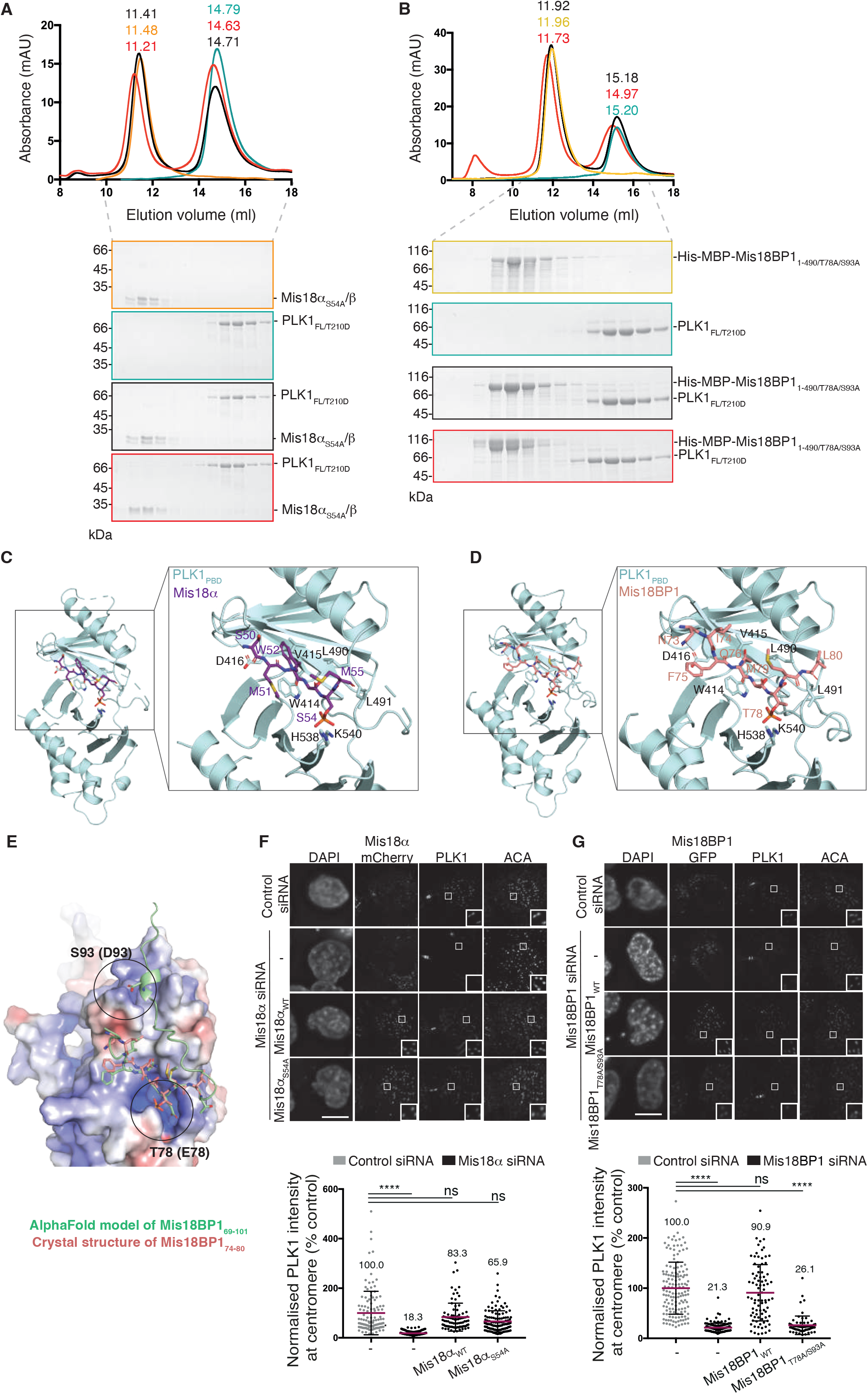
Phosphorylation of Key Residues on Mis18α and Mis18BP1 Mediate Interaction with PLK1 PBD and are Crucial for PLK1 Centromeric Location. (**A**) SEC profiles and corresponding SDS-PAGE analysis of Mis18α_S54A_/β (orange) and PLK1 (green) individually, mixed together (black) and mix together with ATP/Mg^2+^ and incubated to allow phosphorylation (red). (**B**) SEC profiles and corresponding SDS-PAGE analysis of His-MBP-Mis18BP1_1-490/T78A/S93A_ (yellow) and PLK1 (green) individually, mixed together (black) and mix together with ATP/Mg^2+^ and incubated to allow phosphorylation (red). (**C-D**) Crystal structures of PLK1_PBD_ with phosphorylated peptides of (**C**) Mis18α (ASMWSSphM), and (**D**) Mis18BP1 (KNIFQSTphMLTE). The box shows the close-up view of the binding site. PDB: 8S30 and 8S31. (**E**) AlphaFold (*26, 27*) model of PLK1_PBD_ bound to Mis18BP1_69-101_ with phospho-mimic residues are T78E and S93D (highlighted in circles, peptide shown in green) compared with the crystal structure of Mis18BP1_74-80_ with T78ph shown in panel **D** (peptide shown in salmon). (**F-G**) Representative immunofluorescence micrographs and analysis of endogenous PLK1 levels at centromeres in HeLa Kyoto cells during G1 (**F**) when Mis18α was depleted with siRNA oligos and rescued with either Mis18α-mCherry wild-type or Mis18α_S54A_-mCherry and (**G**) when Mis18BP1 was depleted with siRNA oligos and rescued with either Mis18BP1-GFP wild-type or Mis18BP1-GFP_T78A/S93A_. Mean ± SD, n ≥ 85 (**F**) and n ≥ 67 (**G**) from at least 3 independent experiments. Mean values are denoted on graphs. Data were analysed with Kruskal-Wallis with Dunn’s multiple comparisons test. ^****^ P ≤ 0.0001. All scale bars correspond to 10 µm.

To elucidate the structural basis for how PLK1 interacts with Mis18α and Mis18BP1, we determined high resolution crystal structures of the PLK1_PBD_ with phospho-peptides comprising Mis18α_49-55_ (ASMWSSphM, containing phosphorylated S54) and Mis18BP1_72-82_ (KNIFQSTphMLTE containing phosphorylated T78) at 1.9 and 2.1 Å resolution, respectively (Fig. 2C, 2D, S2F & S2G, Supplementary Table 2). Comparison of the Mis18α_S54ph_-PLK1_PBD_ and the Mis18BP1_T78ph_-PLK1_PBD_ structures revealed that both phospho-peptides bound PLK1 in a similar manner and aligned well at residues M/F (−3), S (−1) and Sph/Tph (0) (Fig. 2C & 2D). Further structural analysis showed that the Mis18α-PLK1_PBD_ complex crystal structure has a buried surface area (BSA) of 930 Å^2^, while the BSA for the Mis18BP1_T78ph_-PLK1_PBD_ structure is 1220 Å^2^, suggesting that although Mis18α and Mis18BP1 use the same binding interface in PLK1, Mis18BP1 is likely to bind PLK1 with higher binding affinity as compared with Mis18α. AlphaFold modelling (*26, 27*) of Mis18BP1 fragment containing phosphomimic mutations of T78 and S93 (T78E/S93D) shows that D93 docks at a positively charged region of PLK1_PBD_ close to the canonical phospho-peptide binding pocket, while E78 docks in a similar orientation as T78ph in the crystal structure. The pocket where S93ph docks has recently been described as an evolutionarily conserved cryptic surface involved in substrate discrimination (*28*). This provides a structural basis for how S93ph might further enhance Mis18BP1 interaction with PLK1 (Fig. 2E).

Overall, these structural analyses reveal the interfaces involved in PBD binding to the Mis18 complex subunits. We hypothesised that not only the phosphorylation but also the binding of PLK1 to the Mis18 complex could be important for PLK1 centromere recruitment and CENP-A deposition.

### PLK1-mediated phosphorylation of Mis18BP1 works upstream of Mis18α, in recruiting PLK1 to centromeres

We performed siRNA-rescue assays in HeLa cells to evaluate the role of PLK1-mediated phosphorylation and binding of Mis18α and Mis18BP1 on PLK1 centromere recruitment. We measured endogenous PLK1 levels at centromeres following depletion of either Mis18α or Mis18BP1 using siRNA oligos and rescue with either Mis18α_WT_-mCherry or Mis18α_S54A_-mCherry, and Mis18BP1_WT_-GFP or Mis18BP1_T78A/S93A_-GFP. All Mis18α and Mis18BP1 constructs localised to centromeres in early G1 (Fig. 2F & 2G). Depletion of Mis18α resulted in the loss of endogenous PLK1 at centromeres, which was rescued by Mis18α_WT_-mCherry (Fig. 2F & S2H). Interestingly, a similar level of rescue was observed with the non-phosphorylatable version of Mis18α (Mis18α_S54A_-mCherry). Depletion of Mis18BP1 also showed reduced PLK1 recruitment to centromeres, which was rescued with Mis18BP1_WT_-GFP, but not with Mis18BP1_T78A/S93A_-GFP (Fig. 2G). These observations suggest that while Mis18α S54 phosphorylation is not directly required for PLK1 centromere recruitment, the interaction of Mis18BP1 with PLK1 mediated by phosphorylated T78 and S93 is essential for PLK1 localisation to centromeres. The loss of PLK1 at centromeres due to Mis18α depletion could be explained by the mutual dependency of Mis18α, Mis18β and Mis18BP1 for their centromere localisation (*7*). Our *in vitro* data shows that Mis18α_S54A_ can still form a complex with Mis18BP1 (Fig. S2C). Thus, in the rescue experiment with Mis18α_S54A_-mCherry, Mis18BP1 would still be present at centromeres, available to recruit PLK1.

### PLK1-mediated phosphorylation of Mis18α and Mis18BP1 is required for new CENP-A loading at centromeres

To further dissect the role of PLK1 phosphorylation of Mis18α and Mis18BP1 on new CENP-A incorporation at endogenous centromeres *in vivo*, we performed CENP-A-SNAP deposition assays using a HeLa cell line constitutively expressing SNAP-tagged CENP-A (*19, 29*). CENP-A-SNAP deposition assays utilise a quench-chase-pulse labelling strategy to selectively label newly synthesised CENP-A when it becomes deposited on chromatin (*29*). Mis18α and Mis18BP1 were depleted using siRNA oligos in separate experiments and rescued by transiently expressing either Mis18α_WT_-mCherry or Mis18α_S54A_-mCherry and Mis18BP1_WT_-GFP or Mis18BP1_T78A_-GFP, Mis18BP1_S93A_-GFP, Mis18BP1_T78A/S93A_-GFP. While Mis18α_WT_-mCherry rescued new CENP-A deposition in cells depleted of endogenous Mis18α, Mis18α_S54A_-mCherry showed a significant reduction in new CENP-A loading (Fig. 3A). Expression of Mis18BP1_WT_-GFP in cells depleted of Mis18BP1 rescued new CENP-A loading, while Mis18BP1_T78A_-GFP, Mis18BP1_S93A_-GFP and Mis18BP1_T78A/S93A_-GFP led to reduced new CENP-A deposition, with Mis18BP1_T78A/S93A_-GFP showing the strongest effect (Fig. 3B). Overall, these findings indicate that PLK1 phosphorylation of both Mis18α and Mis18BP1 is essential for new CENP-A loading. Consistent with these results, when TetR-eYFP-Mis18α_WT_ was ectopically tethered to an alphoid^tetO^ array (integrated into a chromosome arm of a HeLa 3-8 cell line (*30*)) in the presence of the PLK1 inhibitor BI2536, we observed a decrease in CENP-A levels at the tethering site as compared with control cells (Fig. S3A).

**Fig. 3.**
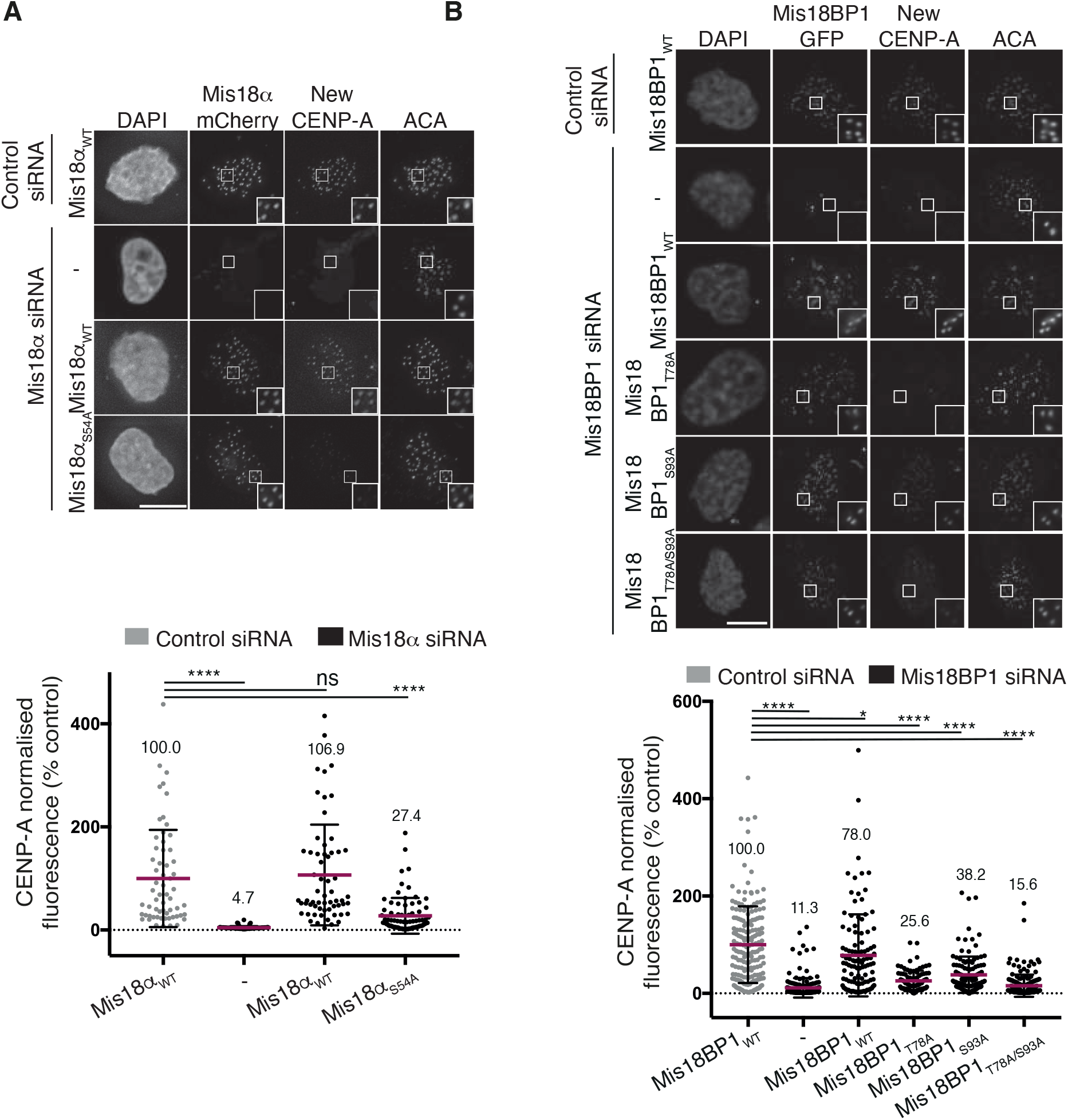
PLK1-mediated Phosphorylation of Key Residues on Mis18α and Mis18BP1 are Required for Proper CENP-A Loading. (**A-B**) Representative immunofluorescence and analysis of new CENP-A-SNAP incorporation at centromeres in a HeLa cell line constitutively expressing SNAP-tagged CENP-A during G1 (**A**) when Mis18α was depleted with siRNA oligos and rescued with either Mis18α-mCherry wild-type or non-phosphorylatable mutant (Mis18α_S54A_-mCherry) or (**B**) when Mis18BP1 was depleted with siRNA oligos and rescued with either Mis18BP1-GFP wild-type or non-phosphorylatable mutants (Mis18BP1-GFP_T78A_, Mis18BP1-GFP_S93A_ or Mis18BP1-GFP_T78A/S93A_). Mean ± SD, n ≥ 61 (**A**) and n ≥ 89 (**B**) from at least 3 independent experiments. Mean values are denoted on graphs. Data were analysed with Kruskal-Wallis followed by Dunn’s multiple comparisons test. ^****^ P ≤ 0.0001, ^*^ P ≤ 0.05. All scale bars correspond to 10 µm.

### PLK1 phosphorylation cascade on the Mis18 complex controls HJURP recruitment to centromeres

To investigate the mechanistic role of PLK1-mediated phosphorylation of the Mis18 complex on new CENP-A loading, we first asked if PLK1 controls new CENP-A deposition by regulating HJURP centromere recruitment. We assessed HJURP levels at endogenous centromeres in G1 cells where either Mis18α or Mis18BP1 was depleted with siRNA oligos and rescued with the wild-type protein or phospho-mutants (either non-phosphorylatable or phosphomimic). We found that Mis18α depletion disrupted HJURP localisation to centromeres, which could be rescued with Mis18α_WT_-mCherry (Fig. 4A), whereas expression of Mis18α_S54A_-mCherry resulted in a significant reduction of HJURP recruitment to centromeres. Likewise, depletion of Mis18BP1 also caused a reduction in HJURP at centromeres which was rescued by expressing Mis18BP1_WT_-GFP, but not by the expressing Mis18BP1_T78A/S93A_-GFP (Fig. 4B). Interestingly, while the expression of Mis18α_S54D_-mCherry led to a significant increase of HJURP levels at centromeres, expression of Mis18BP1_T78D/S93D_ did not show any increase in HJURP centromere recruitment (Fig. 4A-B). These data, together with the findings in Fig. 2F and 2G, suggests that PLK1-mediated phosphorylation of both Mis18α S54 and Mis18BP1 T78 and S93 are required for HJURP centromere recruitment, but while Mis18α S54 directly modulates HJURP recruitment, the Mis18BP1 T78 and S93 phosphorylations work upstream of Mis18α phosphorylation by recruiting PLK1 to centromeres.

**Fig. 4.**
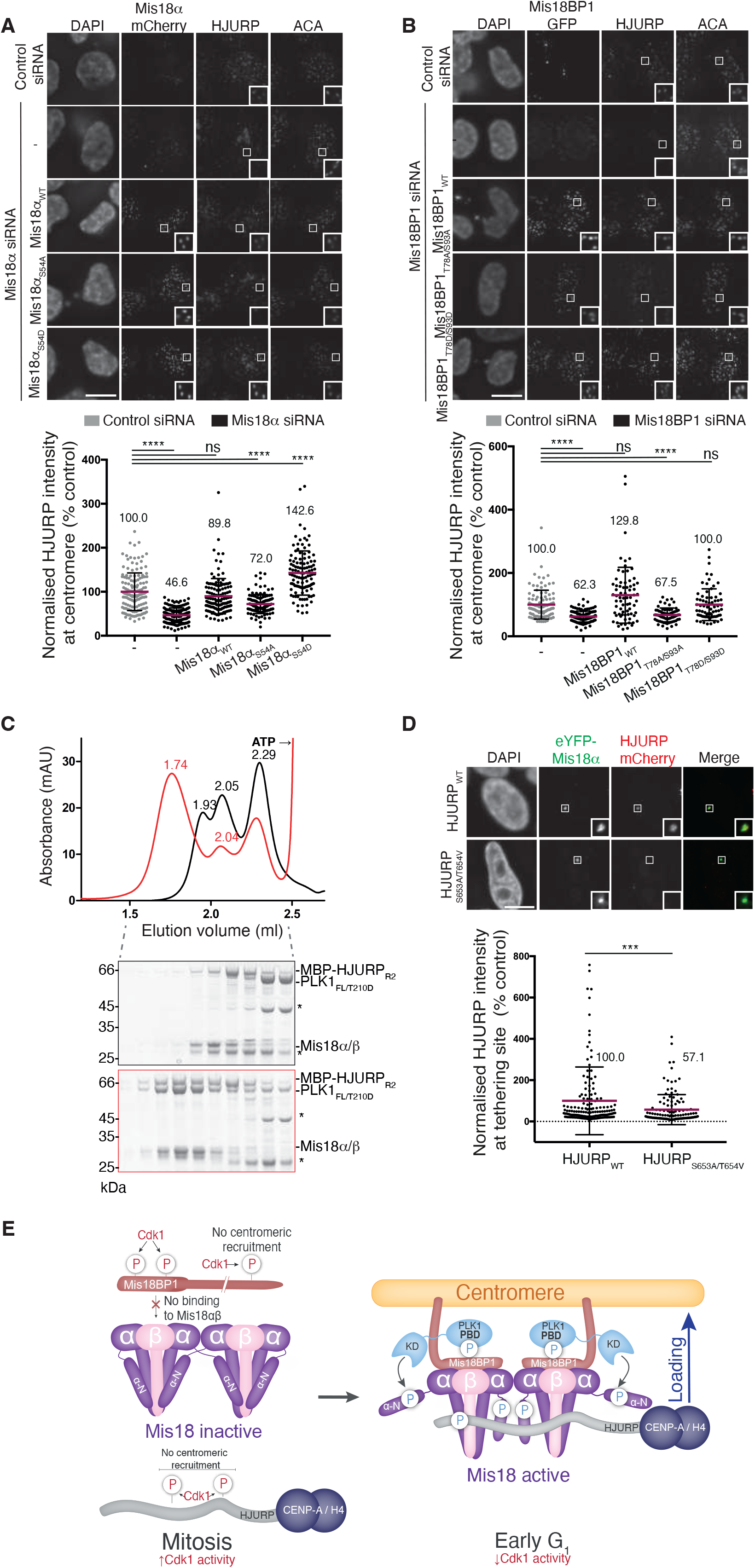
PLK1 Phosphorylation Cascade activates the Mis18 complex to achieve HJURP Centromere Recruitment and CENP-A Deposition. (**A-B**) Representative immunofluorescence micrographs and analysis of endogenous HJURP levels at centromeres in HeLa Kyoto cells during G1 upon (**A**) Mis18α depletion using siRNA and rescue with either Mis18α-mCherry wild-type or phospho-mutants (Mis18α_S54A_-mCherry or Mis18α_S54D_-mCherry), (**B**) Mis18BP1 depletion using siRNA and rescue with either Mis18BP1-GFP wild-type or phospho-mutants (Mis18BP1-GFP_T78A/S93A_ or Mis18BP1-GFP_T78D/S93D_. Mean ± SD, n ≥ 114 (**A**) and n ≥ 67 (**B**) from at least 3 independent experiments. Mean values are denoted on graphs. Data were analysed with Kruskal-Wallis followed by Dunn’s multiple comparisons test. ^****^ P ≤ 0.0001. All scale bars correspond to 10 µm. (C) SEC profiles and corresponding SDS-PAGE analysis of Mis18α/β mixed with PLK1 and His-MBP-HJURP_541-748_ (R2) with no phosphorylation by PLK1 (black) and mixed together with ATP/Mg^2+^ and incubated to allow phosphorylation (red). Asterisks denote contaminations that do not interfere with binding. (**D**) Representative immunofluorescence micrographs and analysis of HJURP-mCherry wild-type or HJURP_S653A/T654V_-mCherry recruitment by TetR-eYFP-Mis18α_WT_ to the alphoid^tetO^ array in HeLa 3-8 cells. Mean ± SD, n ≥ 127 from at least 3 independent experiments. Mean values are denoted on graphs. Data were analysed using a Mann-Whitney U test. ^***^ P ≤ 0.001. All scale bars correspond to 10 µm. (**E**) Mechanistic model proposed describing the role of PLK1 phosphorylation cascade in facilitating HJURP centromere recruitment and CENP-A loading.

### PLK1 phosphorylation cascade activates the Mis18α/β complex to facilitate HJURP binding

It has been previously shown that HJURP interacts with the triple helical bundle of Mis18α/β via its C-terminal HCTD1 and HCTD2 domains (referred to as R1 and R2 domains), and interactions of both domains are essential for CENP-A loading (*24*). Interestingly, HJURP-HCTD2 interaction is possible only when the N-terminal α-helical region of Mis18α is removed, suggesting a regulation involving the Mis18α N-terminal region (*24*). Supporting this notion, our recent structural analysis of the Mis18 complex revealed extensive intramolecular interaction between the Mis18α N-terminal α-helical region and the C-terminal triple helical bundle of Mis18α/β, particularly close to the HJURP contact region described in Pan et al. (*23, 24*). Moreover, our data presented here shows that several PLK1 phosphorylation sites, including the critical Mis18α S54, are located in the N-terminal region of Mis18α (Fig. 1A & Fig. S3B). These observations together led us to conclude that the intramolecular interaction between the Mis18α N-terminal α-helical region keeps the Mis18α/β complex in an inactive/closed conformation occluding HJURP binding. Accordingly, when we tethered TetR-eYFP-Mis18α_54-223_, lacking the first 53 amino acid residues of Mis18α, and assessed its ability to recruit HJURP and load CENP-A to the tethering site using ectopic tethering assays, we observed a two-fold increase of HJURP and CENP-A levels compared to TetR-eYFP-Mis18α_FL_ (Fig. S3C & S3D). These data confirms that the N-terminal region of Mis18α negatively regulates HJURP recruitment and CENP-A deposition. We also observed that the N-terminal α-helical region of Mis18α is required for PLK1 binding (Fig. S3E). Thus, we hypothesised that PLK1 phosphorylation likely regulates Mis18α/β-HJURP interaction by relieving the inactive conformation of Mis18α/β.

In agreement with our hypothesis, we did not observe complex formation when recombinant Mis18α/β complex was mixed with His-MBP-HJURP_541-748_ (R2) and analysed by SEC (Fig. 4C, black panel). However, a robust Mis18α/β/HJURP/PLK1 complex was formed when PLK1 was allowed to phosphorylate Mis18α/β and HJURP (Fig. 4C, red panel). Similar observations were made when we performed the experiments with the Mis18 complex (Mis18α/β/Mis18BP1) (Fig. S4A). Remarkably, PLK1-mediated phosphorylation of HJURP R2 and HJURP R1R2 is more efficient when in the presence of the Mis18α/β complex (Fig. S4B-C). Hence, we wondered if PLK1 phosphorylation of HJURP might also contribute to efficient HJURP-Mis18 complex interaction and CENP-A loading. Amino acid sequence analysis of the C-terminal region of HJURP identified two residues, S653 and T654, within the HJURP R2 domain that could act as possible PLK1 phosphorylation/binding sites (Fig. S4D). AlphaFold modelling (*31*) provided a structural model where a HJURP peptide spanning S653 and a phosphomimic E654 interacts with PLK1_PBD_ in a binding mode similar to that of Mis18α and Mis18BP1 (Fig. S4E). Supporting our hypothesis, ectopic tethering of TetR-eYFP-Mis18α_WT,_ when co-expressed with either HJURP_WT_-mCherry or HJURP_S653A/T654V_-mCherry in the HeLa 3-8 cell line, revealed that the two residues in the predicted PLK1 phospho-sites are needed for efficient HJURP binding to Mis18α as mutating them to non-phosphorylatable amino acid residues reduced Mis18α_WT_ ability to recruit HJURP to the ectopic site (Fig. 4D).

## Discussion

Preserving centromere identity during the cell cycle is of paramount importance. Centromeres act as microtubule attachment sites that harness spindle force to drive chromosome segregation and as sites that hold the sister chromatids together until all chromosomes achieve bi-orientation (*32, 33*). DNA-replication mediated dilution of CENP-A levels poses a threat, as CENP-A levels below a particular threshold will lead to loss of centromere identity. To counter this, the CENP-A loading machinery restores original CENP-A levels by actively depositing correct amounts of CENP-A at centromeres at the right time (*7, 13-15*). This is crucial since incorrect levels and mislocalisation of CENP-A can lead to genomic instability (*5, 34*), whilst unregulated CENP-A deposition at centromeres throughout the cell cycle causes mitotic defects (*19*). However, many questions remain on how the CENP-A loading machinery restores the original levels of CENP-A at a specific site during a defined time frame in a DNA sequence-independent manner. In recent years, we have started to gain mechanistic insights into this process.

Several licensing steps involving CDKs and PLK1 have been suggested to achieve the spatiotemporal control of CENP-A deposition. Thus far, we know that both the Mis18 complex and HJURP are regulated by CDKs, with the Mis18 complex additionally regulated by PLK1 (*16-20*). CDKs negatively regulate the Mis18 complex formation required for HJURP centromere recruitment, through the phosphorylation of Mis18BP1 residues T40 and S110 (*16, 17*). Our recent structural analysis of the Mis18 complex revealed that these phosphorylation sites on Mis18BP1 lie at its binding interface with the Mis18α/β complex, providing the structural basis for this regulation (*23*). CDKs have also been shown to control Mis18BP1 centromere localisation through the phosphorylation of Mis18BP1 T653, which is likely to perturb Mis18BP1 interaction with CCAN (*8, 9, 12, 20*). These phosphorylation events together inhibit CENP-A deposition until late mitosis/G1 when CDK activity decreases. Unlike CDKs activity, PLK1 is recruited to the centromere during G1 in a Mis18 complex-dependent manner, and its activity promotes Mis18 complex centromere localisation and subsequent CENP-A loading, through the phosphorylation of amino acid residues within the N-terminal half of Mis18BP1 (Mis18BP1_1-490_, a fragment capable of associating with the centromere) (*19*). However, a mechanistic understanding of how this key licensing process is established, what are the crucial molecular events constituting this licensing step and how these translate into the regulation of CENP-A deposition remains unclear.

In this study, we utilised biochemical, structural, and *in vivo* functional methods to dissect the PLK1-mediated licensing mechanism of CENP-A deposition. We: (i) reveal that amino acid residues of Mis18BP1 (T78 and S93) and Mis18α (S54), upon self-priming phosphorylation by PLK1, act as docking sites for PLK1_PBD_; (ii) show, by determining high-resolution crystal structures and using AI-based structural modeling, that Mis18BP1 binding by PLK1_PBD_ exploits both phosphorylated T78 (forming a canonical PLK1_PBD_ binding motif) and phosphorylated S93, where the latter engages in a pocket adjacent to the canonical PLK1_PBD_ phospho-peptide binding pocket; (iii) show that PLK1 phosphorylation and binding site on Mis18α lies within the helical region which we have previously shown to make intramolecular interaction with the HJURP binding site of the Mis18α/β complex; (iv) demonstrate that PLK1 phosphorylation/binding of Mis18α_S54_ activates Mis18α/β, making it compatible for HJURP binding, by relieving the intramolecular interaction between Mis18α N-terminal helical region and HJURP binding surface; and (v) show that phosphorylation of Mis18BP1 and binding of PLK1 works upstream of the Mis18α/β phosphorylation, and that PLK1 binding and phosphorylation of HJURP might further enhance Mis18α/β-HJURP interaction and facilitate HJURP centromere recruitment.

Taken all together, we provide a mechanistic model in which PLK1 establishes a phosphorylation cascade (Fig. 4E). It starts with PLK1 phosphorylation of centromere associated Mis18BP1 at G1, which then provides a docking site for PLK1. The Mis18BP1 bound PLK1 then phosphorylates and interacts with Mis18α/β. This achieves two things: concentrating PLK1 at the centromere at the right time and, most importantly, relieving the intramolecular inhibition of the Mis18α/β complex for robust HJURP binding. The HJURP binding by the Mis18α/β appears to facilitate PLK1 phosphorylation of HJURP, which is likely to contribute further to the centromere recruitment of HJURP. Perturbing the PLK1 docking site on Mis18BP1 (T78A and S93A) while abolishing PLK1 centromere recruitment, did not majorly affect the centromere localisation of the Mis18 complex. This emphasises that the PLK1 licensing role is not just regulating the centromere association of the Mis18 complex as previously thought (*19*), but activating the Mis18α/β complex by making it compatible for efficient HJURP binding. Our mechanistic model also explains why artificial centromere targeting of Mis18BP1 alone is not sufficient to bypass the requirement of PLK1 activity as observed elsewhere (*19*). In the future, it would be interesting to investigate if PLK1 has any role in the downstream maturation process that stably incorporates CENP-A nucleosomes into the centromeric chromatin (*35, 36*). Notably, the CENP-A loading machinery and CCAN components are suggested to enrich at centromeres during S-phase to ensure CENP-A nucleosome inheritance during DNA replication (*37, 38*). Our work also paves the way for further exciting questions on whether a similar phosphorylation cascade acts on the CENP-A loading machinery to warrant accurate inheritance of CENP-A nucleosomes during DNA replication. The findings and conclusions of this work are broadly in agreement with the work of Conti *et al*. (ref pending).

## Materials and Methods

### Plasmids

Codon optimised (GeneArt) Mis18α and Mis18β genes were cloned into expression vectors pET His6 TEV (9B) and pET His6 msfGFP TEV (9GFP, Addgene plasmids #48284 and #48287, a gift from Scott Gradia), respectively, and combined to form a polycistronic vector. Mis18BP1_1-490_ was cloned from a codon optimised sequence (GeneArt) into the pET His6 MBP TEV (14C, Addgene plasmid #48309, a gift from Scott Gradia). Mis18BP1_20-130_ was cloned into pEC-K-3C-His-GST. Codon optimised HJURP_541-748_ (GeneArt) fragments were cloned into pET His6 MBP TEV (14C). PLK1_FL/T210D_ and PLK1_370-603_ (called PLK1_PBD_ in this study) were cloned into the pEC-A-HI-SUMO expression vector.

For cell studies, non-codon optimised sequences for Mis18α, Mis18BP1 and HJURP were cloned into pcDNA3 mCherry and pcDNA3 GFP vectors (6B and 6D, Addgene plasmids #30125 and #30127, a gift from Scott Gradia). Mis18α was also cloned into TetR-eYFP-IRES-Puro vector. All mutations were generated using the Quikchange site-directed mutagenesis method (Stratagene).

### Expression and recombinant protein purification

A polycistronic vector containing genes for full-length His-tagged Mis18α and full-length His-GFP-tagged Mis18β, His-tagged Mis18α_54-223_ and full-length His-GFP-tagged Mis18β, His-SUMO-PLK1_FL/T210D_, His-SUMO-PLK1_PBD_, His-GST-Mis18BP1_20-130_ and His-MBP-HJURP_541-748_ were used to express in *E. coli* BL21 Gold (DE3) whilst His-MBP-Mis18BP1_1-490_ was expressed in pLysS (DE3). Cultures were grown in LB (Mis18α/β, Mis18BP1_20-130_ and HJURP_541-748_) or super broth media (PLK1_FL/T210D_, PLK1_PBD_ and Mis18BP1_1-490_) at 37°C to O.D. of 0.6-1.0 and the temperature was reduced to 18°C for an 1 h and cultures induced with 0.35 mM IPTG overnight.

Cells were lysed by sonication in lysis buffer (Mis18α/β: 20 mM Tris, pH 8.0, 250 mM NaCl, 35 mM Imidazole and 2 mM β-ME; Mis18BP1_20-130_: 20 mM Tris, pH 8.0, 500 mM NaCl, 35 mM Imidazole and 2 mM β-ME; Mis18BP1_1-490_: 20 mM potassium, phosphate, pH 7.4, 100 mM NaCl, 35 mM Imidazole, and 2 mM β-ME; PLK1_FL/T210D_: 50 mM MOPS, pH 7.5, 350 mM NaCl, 35 mM imidazole, and 2 mM β-ME; PLK1_PBD_: 20 mM Tris, pH 8.0, 500 mM NaCl, 35 mM Imidazole, 2 mM β-ME and HJURP_541-748_: 20 mM HEPES, pH 7.5, 3000 mM NaCl, 35 mM Imidazole and 2 mM β-ME) and supplemented with 1 mM PMSF, 10 µg/ml DNase, 5 mM MgCl_2_ and cOmplete (EDTA-free, Sigma) and purified using HisTrap™ HP 5 ml column (Cytiva). The protein-bound resin was washed with lysis buffer, followed by a chaperone buffer (lysis buffer containing 1 M NaCl, 50 mM KCl, 10 mM MgCl_2_ and 2 mM ATP, except for PLK1_FL/T210D_, which just had 1 M NaCl). No chaperone wash was used for PLK1_PBD_. After an additional lysis buffer wash, proteins were eluted with lysis buffer containing 400 mM imidazole. Proteins were then dilysed overnight into dialysis buffer (Mis18α/β: 20 mM Tris pH 8.0, 150 mM NaCl and 2 mM DTT; Mis18BP1_20-130_: 20 mM Tris, pH 8.0, 100 mM NaCl and 2 mM DTT; Mis18BP1_1-490_: 20 mM Tris, pH 7.5, 100 mM NaCl and 2 mM DTT; PLK1_FL/T210D_: 50mM MOPS, pH 7.5, 200 mM NaCl and 2 mM DTT and PLK1_PBD_: 20 mM Tris, pH 8.0, 500 mM NaCl, and 2 mM DTT) and cleaved with either TEV or SENP2 proteases as required.

Except for PLK1_FL/T210D,_ PLK1_PBD_ and HJURP_541-748_, all other proteins were further purified by anion exchange chromatography using HiTrap™ Q HP (Cytiva), the relevant fractions were then pooled and concentrated. Proteins were the injected onto Superdex® 75 Increase 10/300 GL (Mis18BP1_20-130_ and PLK1_PBD_), Superdex® 200 Increase 10/300 GL or Superose® 6 10/300 GL (Mis18α/β/Mis18BP1_1-490_) column equilibrated with SEC buffer (Mis18α/β: 20 mM Tris. pH 8.0, 250 mM NaCl and 2 mM DTT; Mis18BP1_20-130_: 20 mM Tris, pH 8.0, 100 mM NaCl, and 2 mM DTT; Mis18BP1_1-490_: 20 mM Tris, pH 7.5, 200 mM NaCl and 2 mM DTT; Mis18α/β/Mis18BP1_1-490_: 20 mM Tris. pH 8.0, 350 mM NaCl and 2 mM DTT; PLK1_FL/T210D_: 50 mM MOPS, pH 7.5, 150 mM NaCl and 2 mM DTT; PLK1_PBD_: 20 mM Tris, pH 8.0, 500 mM NaCl, 2 mM DTT and HJURP_541-748_: 20 mM HEPES, pH 7.5, 3000 mM NaCl, 2 mM DTT). Fractions were analysed on SDS-PAGE stained with Coomassie blue.

### Protein interaction trials

All proteins were phosphorylated by the addition of 2- or 3-mM ATP and 10 mM MgCl_2_ before incubated at 33°C for 45 min at 500 rpm. For initial interaction, the following conditions were used: Mis18α/β-PLK1_FL/T210D_ and Mis18α/β-PLK1_PBD_ interactions were performed using a Superdex® 200 Increase 10/300 GL column (Cytiva) was equilibrated with a buffer containing 50 mM MOPS, pH 7.5, 150 mM NaCl, and 2 mM DTT. For His-MBP-Mis18BP1-PLK1_FL/T210D_ interactions, Superdex® 200 Increase 10/300 GL column was equilibrated with a buffer containing 50 mM MOPS, pH 7.5, 350 mM NaCl, and 2 mM DTT. For Mis18α/β/Mis18BP1-PLK1_FL/T210D_ interactions, Superose® 6 10/300 GL column (Cytiva) was equilibrated with a buffer containing 50 mM MOPS, pH 7.5, 150 or 350 mM NaCl, and 2 mM DTT. Subsequent SEC was performed with Superose® 6 5/150 column (Cytiva) equilibrated with 50 mM MOPS, pH 7.5, 150 mM NaCl, and 2 mM TCEP. For each interaction trial, samples contained identical protein molarities and sample volumes were used.

Amylose pulldown assay has been described previously (*24*). In brief, 5 mM His-MBP-Mis18BP1_1-490_ was mixed proteins indicated and with 0.5-10mM PLK1 as specified (with or without 2 mM ATP and 10 mM MgCl_2_) and incubated at 33ºC for 45 mins at 500 rpm. To check PLK1_PBD_ binding, 5 mM PLK1_PBD_ was added after incubation. Sample was diluted in a buffer containing 20 mM HEPES, pH 8.0, 150 mM NaCl, 1 mM TCEP, and 0.01% Tween-20 to make up a total volume of 40 µl. 8 µl (25%) sample was taken as input and the rest of the samples incubated with 50 µl amylose resin (Thermo Fisher Scientific) which had been washed in buffer before incubating for 90 minutes at 4ºC in a rotating mixer. Beads were then washed with 500 µl 5-6 times with either lysis buffer (His-MBP-Mis18BP1_1-490_ with PLK1_PBD_) or lysis buffer containing 350 mM NaCl (His-MBP-Mis18BP1_1-490_ and Mis18 complex with PLK1) and protein eluted in SDS-PAGE loading dye by boiling at 95ºC for 5 minutes. Input and bound samples were analysed on 12% SDS-PAGE gels stained using Coomassie Blue.

### Mass spectrometry (MS)

Phosphorylated samples of interest were purified by SEC to ensure homogeneity, then were run on NuPAGE™ 4-12% Bis-Tris (Invitrogen) pre-cast gels and stained using InstantBlue™ Coomassie Stain (Expedeon). During sample preparation, bands of interest were excised and reduced using 10 mM DTT at 37ºC for 30 mins and alkylated with 55 mM iodoacetamide for 20 mins at room temperature. Trypsin buffer containing 13 ng/mL trypsin (Promega) in 10 mM ammonium bicarbonate and 10% (v/v) acetonitrile was then added and incubating overnight at 37ºC. The peptides were then loaded onto C18-StageTips (*39*).

LC-MS/MS analysis was performed using Orbitrap Fusion Lumos (Thermo Fisher Scientific). The peptide separation was carried out on an EASY-Spray column (Thermo Fisher Scientific). Mobile phase A consisting of water and 0.1% (v/v) formic acid and Mobile phase B consisting of 80% (v/v) acetonitrile and 0.1% (v/v) formic acid were used. The digested peptides were loaded at a flow rate of 0.3 ml/min and eluted at a flow rate of 0. 2ml/min using a linear gradient of 2% to 40% mobile phase B over 55 mins, followed by 40% to 95% mobile phase B increase over 11 mins. The eluted peptides were then added to the mass spectrometer and their data acquired in a data-dependent mode with a 3 second acquisition cycle. The Orbitrap was used to record the precursor spectra with a resolution of 120,000. The ions with precursor charges in the range of 3+ to 7+ were fragmented with a collision energy of 30 using high-energy collision dissociation (HCD) and their fragmentation spectra were recorded in the Orbitrap with a resolution of 30,000. Raw files containing mass spectrometric data were processed using MaxQuant 1.6.1.0 (*40*).

### Crystallisation, data collection, and structure determination

A custom peptide was designed and ordered from Peptide Synthetics containing the following sequence: ASMWSSphM (S54ph peptide), solubilised in 15% isopropanol and 75% DMSO and KNIFQSTphMLTE (T78ph peptide), solubilised in 100% DMSO.

For Mis18α-PLK1_PBD_ crystal structure, PLK1_PBD_ was concentrated to 6 mg/ml and mixed with two times molar excess of S54ph peptide The mixture was incubated on ice for at least 1 h before setting up crystallisation trays with Morpheus® (Molecular Dimensions) screen using the ART Robbins Crystal Gryphon crystallisation robot in 96-well sitting drop MRC plates at 18ºC. Morpheus plate containing 0.09M Halogen mix (NaF, NaBr, NaI), 0.1 M buffer system 2 (Sodium HEPES and MOPS, pH7.5), 37% precipitant mix MPD_P1K_P3350 (MPD (racemic), PEG 1K, PEG 3350). The crystals were frozen in liquid nitrogen and sent for data collection to Diamond Light Source beamline i04 (Oxford, United Kingdom).

For Mis18BP1-PLK1_PBD_ structure, PLK1_PBD_ was purified and concentrated to 12 mg/lL and mixed with two times molar excess of T78ph peptide. The mixture was incubated on ice for over 1 h before setting up crystallisation trays with homemade screens using the ART Robbins Crystal Gryphon crystallisation robot in 96-well sitting drop MRC plates at 18ºC. Crystals were obtained in the condition 50 mM MES, pH 6.0, 20 mM Sodium Oxalate, 1.2 M Sodium Malonate, then frozen in liquid nitrogen and sent for data collection to Diamond Light Source beamline i24 (Oxford, United Kingdom).

Crystal structures were resolved by molecular replacement with PHASER (*41*). The coordinates of the structure of PLK1_PBD_ (RCSB PDB ID: 5NFU; model used without the bound peptide) were used as a template for molecular replacement. PHENIX suite was then used to perform subsequent rounds of refinements of the structures until a clear density for the peptide was found (*42*). Iterative rounds of model building and structural superpositions were then performed using COOT (*43*).

### Structure modelling

Structural models for PLK1_PBD_ bound to phosphomimic peptides of Mis18BP1 and HJURP were generated using the AlphaFold (*26, 27*). AlphaFold multimer installed locally was used to generate the PLK1_PBD_ -Mis18BP1 structure, while ColabFold AlphaFold2 (*31*) available on google colab was used for the PLK1_PBD_ – HJURP structure. Predicted structures were analysed and figures were generated using PyMOL (*44*).

### Western blot

To study the expression levels of Mis18α and each of the Mis18α-mCherry constructs, HeLa CENP-A SNAP cells were transfected in 12-well dishes as described above and solubilised 1× SDS-PAGE loading dye, boiled for 5 min, and analysed by SDS-PAGE followed by Western blotting. The antibodies used for the immunoblot were rabbit anti-tubulin (1:10,000; ab18251; Abcam) and mouse anti-Mis18α (1:100; 25G8, Helmholtz Zentrum München). Secondary antibodies used were goat anti-mouse 680 and donkey anti-rabbit 800 (1:5,000, LI-COR). Immunoblots were imaged using the Odyssey CLx system.

### Cell culture, immunofluorescence, and quantification

Mammalian cells were maintained in DMEM (Gibco) supplemented with 10% FBS (Biowest) and penicillin/streptomycin (Gibco) and incubated at 37ºC in a 5% CO_2_ incubator.

CENP-A-SNAP assay was performed as described previously (*29*), using HeLa CENP-A-SNAP expressing cKM58 cell line (a gift from Iain Cheeseman (*19*)). Cells were grown on coverslips in a 12-well plate and allowed to grow for ∼16 h. For Mis18α, deletion of endogenous protein using siRNA (4392420-s28851, ThermoFisher Scientific) and rescue experiments were performed using jetPRIME® (Polyplus Transfections) according to the manufacturer’s instructions. For Mis18BP1, two steps of transfections were performed: DNA using XtremeGENE™ 9 (Roche) followed by siRNA (4392420-s30722, ThermoFisher Scientific) transfection using jetPRIME® following the manufacturer’s instructions. AllStar negative control siRNA (1027280, Qiagen) was used in both experiments. Cells were transfected with 200 ng DNA for Mis18α constructs and 600 ng of DNA for Mis18BP1 constructs, and 2.5 ul of 10mM siRNA oligos. The following day after DNA transfection, 1 mM thymidine was added to the cells and incubated for 19 h. Thymidine was then removed by washing cells with culture media and blocking of existing CENP-A was performed by treatment with 10 mM SNAP-Cell® Block BTP (S9106S, NEB) for 30 min. Cells were thoroughly washed with culture media to get rid of unbound BTP, then a second wash performed after 30 min. 4 h after the initial thymidine release, the cells were incubated with 1 µM S-trityl-L-cysteine (STLC) for another 15 h before releasing by washing with media. 2 h after STLC release, newly deposited CENP-A was labelled with 3 µM SNAP-Cell® 647-siR (S910102S, NEB) for 30 mins before washing excess with media and allowing to grow for a further 30 mins. Cells grown on coverslips were pre-extracted with 0.1% triton in 1X PBS (only for Mis18α) and fixed with 4% paraformaldehyde (PFA, in 1X PBS) for 10 mins at 37ºC (room temperature for Mis18BP1). The antibodies used for indirect immunofluorescence anti-ACA (1:300 dilution, 15-235, Antibodies Inc.), Alexa Fluor® 488 donkey anti-human (1:300 dilution, 709-546-149, Jackson Immunoresearch) secondary antibody for Mis18α and goat anti-human Rhodamine (1:200 dilution, 109-025-003, Immunoresearch) secondary antibody for Mis18BP1. Coverslips were mounted on glass slides using Vectashield® anti-fade mounting medium with DAPI staining (Vector Laboratories).

To assessed endogenous levels of HJURP and PLK1, HeLa Kyoto cells were grown and transfected as stated above. The day after transfection cells were synchronised with 1 µM STLC for 15 h, then released for 2 h. For HJURP immunostaining, cells were pre-extracted with 0.1% triton, then fixed with 4% PFA whilst PLK1 cells were fixed with methanol. The following antibodies were used for indirect immunofluorescence: anti-ACA (1:300, 15-235, Antibodies Inc.), anti-HJURP (1:200, HPA008436, Atlas Antibodies) and anti-PLK1 (1:500, ab17057, Abcam). Secondary antibodies used were donkey anti-rabbit FITC, goat anti-rabbit TRITC, donkey anti-mouse FITC, donkey anti-mouse TRITC and donkey anti-human Cy5 (1:300, 711-095-152, 111-025-006, 715-025-150, 715-095-150, 709-175-149, Jackson Immunoresearch). Coverslips were mounted on glass slides using Vectashield® anti-fade mounting medium with DAPI staining.

The HeLa 3-8 cell line containing a synthetic a-satellite (alphoid) DNA array integration with tetO sites (alphoidtetO array) integrated in a chromosome arm was used for tethering experiments (*30*). To assess CENP-A deposition at the tethering site, 500 ng of TetR-eYFP-Mis18α vectors were transfected using Opti-MEM (Invitrogen) and XtremeGene-9 (Sigma) following manufacturer’s instructions. For HJURP recruitment at the tethering site, 1 µg of tetR-eYFP-Mis18α and pcDNA3-mCherry-HJURP were used and incubated for 48 h. Where indicated, cells were treated with 100 nM of BI2536 (B3200, LKT Laboratories) for 18 h. For HJURP analysis, cells were pre-extracted with 0.5% triton and fixed in 4% PFA. For CENP-A analysis, cells were fixed with methanol and immunofluorescence performed with anti-ACA and donkey anti-human TRITC. Coverslips were mounted on glass slides using Vectashield® anti-fade mounting medium with DAPI staining.

Cells were imaged using Nikon Ti2 Live Imaging Microscope (Nikon) with CFI Plan Apochromat TIRF 100x objective with oil immersion (refractive index = 1.514) using Nikon Elements 5.1 software. The 0.2 µm spaced z-stacks were deconvolved using Huygens (Scientific Volume Imaging) software. Intensities of newly deposited CENP-A-SNAP at endogenous centromeres were then quantified using an automatic custom-made macro (modified from (*45*), zenodo: 10623895) in ImageJ software (NIH, Bethesda). ACA signals were used as reference channels to determine the location of centromeres in a 7×7 pixel box. CENP-A intensity (data channel) was measured in transfected cells and mean signalling intensities were obtained by subtracting the minimum intensities in the square area. Average intensities of each cell were obtained, and fluorescence was normalised percent against the control.

To analyse the intensity of either PLK1 or HJURP at endogenous centromeres an ImageJ plugin was used (zenodo: 10623895). The plugin detects centromeres using the reference channel (ACA) and quantifies mean intensity levels in two other channels to measure expression levels of the transfected vector and either PLK1 or HJURP levels. To quantify the levels of CENP-A or HJURP found at the tethering site, an ImageJ plugin was used (zenodo: 10650818). The plugin detects the point with the highest intensity in the channel with the tethering site, draws a 7-pixel circle around it and detects the mean intensity levels for another channel in the same area.

For each experiment, a minimum of three biological replicates were performed to plot the graph in Prism 7.0 software. Mann Whitney U test or Kruskal-Wallis followed by Dunn’s test were performed in Prism to measure the statistical significance of the obtained results. Shown images are maximum-intensity projections.

## Acknowledgments

We would like to thank the Centre for Optical Instrumentation Laboratory for their help with microscopy and analysis. We would also like to acknowledge Diamond Light Source, where the crystal structure data was collected. In addition, we would like to thank Andrea Musacchio and Duccio Conti for discussion and sharing of unpublished data.

## Funding

Research in AAJ was supported by Wellcome Senior Research Fellowship (202811). AAJ and his team are co-funded by the European Union (ERC, CHROMSEG, 101054950) and the Medical Research Council (MRC, United Kingdom; MR/X001245/1). Views and opinions expressed are however those of the author(s) only and do not necessarily reflect those of the European Union or the European Research Council. Neither the European Union nor the granting authority can be held responsible for them. The Wellcome Centre for Cell Biology is supported by core funding from the Wellcome Trust (203149). P.P. is funded by the Darwin Trust of Edinburgh.

## Authors contributions

Conceptualisation: A.A.J.

Methodology: P.P., B.M-P., A.A., P.P.S, R.T., D.A.K, T.M., A.A.J.

Investigation: P.P., B.M-P., A.A., P.P.S., R.T., J.Z., D.G, V.D.

Funding acquisition: J.R., A.A.J.

Writing-original draft: P.P., B.M-P., A.A., A.A.J.

Writing-review & editing: P.P., B.M-P., A.A., P.P.S., A.A.J.

## Comping interests

Authors declare that they have no competing interests.

## Data and material availability

Crystal structures are deposited in Protein Data Bank (PDB: http://www.rcsb.org/) under the following accession numbers: 8S30 and 8S31. Plugins used to analyse CENP-A-SNAP data, CENP-A_SNAP_2024, and levels at endogenous centromeres, EndogenousCentromeres_Intensity, are deposited in zenodo: 10623895. The plugin used to analyse tethering data, Spot_Intensity, deposited in zenodo: 10650818. All data are available in the main text or the supplementary materials.

## Supplementary

### Supplementary Figure Legends

**Fig. S1. Mis18α/β/Mis18BP1 Interact with PKL1 in a Phospho-Dependent Manner through PLK1**_**PBD**_. (**A**) SEC of His-Mis18α/His-GFP-Mis18β/His-MBP-Mis18BP1_1-490_ (orange) and PLK1 (green) individually, mixed together (black) and mix together with ATP/Mg^2+^ and incubated to allow phosphorylation (red). Black dotted line indicated the void sample run on the SDS PAGE. Asterisk denotes contaminant from the PLK1 purification. (**B**) SEC of Mis18α/β (orange) and SUMO-PLK1_PBD_ (green) individually, Mis18α/β mixed together with sub-stoichiometric amounts of PLK1 (black), Mis18α/β, SUMO-PLK1_PBD_ with sub-stoichiometric amounts of PLK1 (salmon), Mis18α/β mixed together with sub-stoichiometric amounts of PLK1 and ATP/Mg^2+^ then incubated to allow phosphorylation (red), Mis18α/β and SUMO-PLK1_PBD_ mixed together with sub-stoichiometric amounts of PLK1 and ATP/Mg^2+^ then incubated to allow phosphorylation (blue).**(**C-D**) Amylose binding assays to assess the ability of (**C**) His-MBP-Mis18BP1_1-490_ and (**D**) His-Mis18α/His-GFP-Mis18β/His-MBP-Mis18BP1_1-490_ to interact with PLK1_PBD_ when not phosphorylated, phosphorylated with sub-stoichiometric amounts of PLK1 and stoichiometric amounts of PLK1. The left panel shows inputs, and the right panel shows bead-bound fractions.

**Fig. S2. Phosphorylation of Key Residues on Mis18α and Mis18BP1 Mediate Interaction with PKL1**_**PBD**_. (**A**) SEC profiles and corresponding SDS-PAGE analysis of Mis18α_4A_/β (orange, mutations S53A/S54A/S56A/S60A) and PLK1 (green) individually, mixed together (black) and mixed together with ATP/Mg^2+^ and incubated to allow phosphorylation (red). Asterisk denotes contaminant from the PLK1 purification. (**B**) SDS-PAGE analysis of amylose pull-down assays to assess the ability of His-MBP-Mis18BP1_1-490_ wild-type and mutant proteins to interact with PLK1 when not phosphorylated and phosphorylated with stoichiometric amounts of PLK1. The left panel shows inputs, and the right panel shows amylose bead-bound fractions. (**C-E**) SEC profiles and corresponding SDS-PAGE analysis of either (**C**) His-Mis18α_S54A_/His-GFP-Mis18β/His-MBP-Mis18BP1_1-490/T78A/S93A_ or (**D-E**) His-Mis18α/His-GFP-Mis18β/Mis18BP1_1-490/T78A/S93A_ (orange) and PLK1 (green) individually, mixed together (black) and mixed together with ATP/Mg^2+^ and incubated to allow phosphorylation (red) in buffer containing (**C-D**) 150 mM NaCl and (**E**) 350 mM NaCl. Black dotted lines indicate the void sample run on the SDS PAGE. Asterisk denotes contaminant from the PLK1 purification. (**F-G**) Crystal structures of PLK1_PBD_ with phosphorylated peptides displaying 2*F*_o_–*F*_c_ electron density maps for (**F**) Mis18α and (**G**) Mis18BP1. (**H**) Western blots probed using anti-Mis18α and anti-tubulin antibodies showing the transient expression of Mis18α-mCherry when depleted with control or Mis18α siRNA oligos and the level of depletion of endogenous Mis18α by siRNA oligos.

**Fig. S3. PLK1 Activates the Mis18α/β complex by relieving the inhibitory role of Mis18α N-terminal** α**-helical Region**. (**A**) Representative immunofluorescence micrographs and analysis of CENP-A loading at the tethering site via TetR-eYFP-Mis18α in HeLa 3-8 cells during G1 with and without treatment with the PLK1 inhibitor BI2536. Mean ± SD, n ≥ 209 from at least 3 independent experiments. Mean values are denoted on graphs. Data were analysed using a Mann-Whitney U test. ^****^ P ≤ 0.0001. All scale bars correspond to 10 µm. (**B**) AlphaFold model (*23, 26, 27*) of Mis18α (purple), Mis18β (pink) and Mis18BP1 (salmon) where the N-terminal region of Mis18α (turquoise) had been modelled. Grey residues denote HJURP contact regions identified by (*24*). Red arrows highlight the location of Mis18α residue S54. (**C-D**) Representative immunofluorescence micrographs and analysis of the alphoid^tetO^ array in cells expressing TetR-eYFP-Mis18α_FL_ and TetR-eYFP-Mis18α_54-223_ to assess (**C**) recruitment of endogenous CENP-A to the ectopic site in HeLa 3-8 cells and (**D**) recruitment of HJURP-mCherry to the ectopic site in HeLa 3-8 CENP-A SNAP cells. Mean ± SD, n ≥ 48 (**C**) and n ≥ 49 (**D**) from at least 3 independent experiments. Mean values are denoted on graphs. Data were analysed using a Mann-Whitney U test. ^****^ P ≤ 0.0001, ^*^ P ≤ 0.05. All scale bars correspond to 10 µm. (**E**) SEC profiles and corresponding SDS-PAGE analysis of Mis18α_54-223_/β (orange) and PLK1 (green) individually, mixed together (black) and mixed together with ATP/Mg^2+^ and incubated to allow phosphorylation (red).

**Fig. S4. PLK1 Phosphorylation Cascade on Mis18 Complex and HJURP facilities robust Mis18 complex-HJURP Interaction**. (**A**) SEC profiles and corresponding SDS-PAGE analysis of Mis18α/β mixed with Mis18BP1_20-130_, His-MBP-HJURP_541-748_ (R2) and PLK1 with no phosphorylation (black) and mixed together with ATP/Mg^2+^ and incubated to allow phosphorylation by PLK1 (red). (**B-C)** SDS-PAGE analysis of non-phosphorylated and phosphorylated samples of (**B**) Mis18α/β and PLK1, PLK1 and His-MBP-HJURP_541-748_ (R2) and Mis18α/β, PLK1 and His-MBP-HJURP_541-748_ (R2) with sub-stoichiometric and stoichiometric amounts of Mis18α/β. (**C**) Shows the same experiment as in **B** conducted with His-MBP-HJURP_388-748_ (R1R2). (**D**) Multiple sequence alignment for HJURP using MUSCLE (*46*) visualised with Jalview (*47*) with sequences from *Homo sapiens* (*hs*), *Pan troglodytes* (*pt*), *Bos taurus* (*bt*), *Mus musculus* (*mm*) and *Rattus norvegicus* (*rn*). Secondary structure prediction was performed using JPred Secondary Structure Prediction (*48*). Grey dots indicate potential phosphorylated sites, black lines indicate potential PLK1_PBD_ binding sites. (**E**) AlphaFold modelled structure of PLK1_PBD_ with HJURP generated using ColabFold (*31*).

**Supplementary Table 1. Phosphorylated Peptides**. List of all phosphorylated peptides identified via mass spectrometry in Mis18α/β, Mis18BP1_1-490_ and Mis18α/β/Mis18BP1_1-490_ samples phosphorylated by PLK1.

**Supplementary Table 2. Data Collection and Refinement Statistics**.

## Notes

### Competing Interest Statement

The authors have declared no competing interest.

