## Supplementary figures and images for "PLK1-Mediated Phosphorylation Cascade Activates the Mis18 Complex to Ensure Centromere Inheritance"

### Supplemental Figure 1

#### Supplementary 1

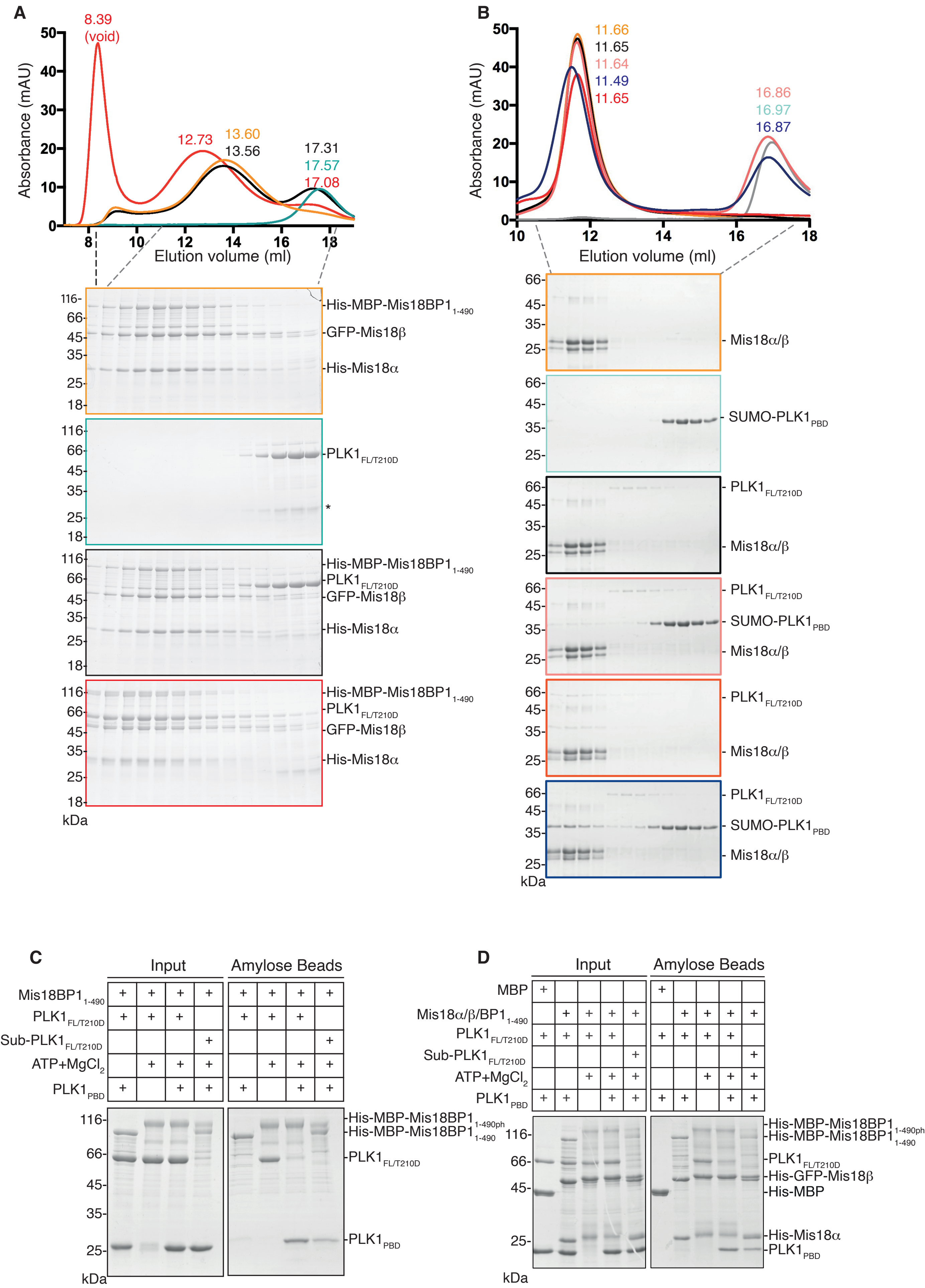

### Supplemental Figure 2

#### Supplementary 2

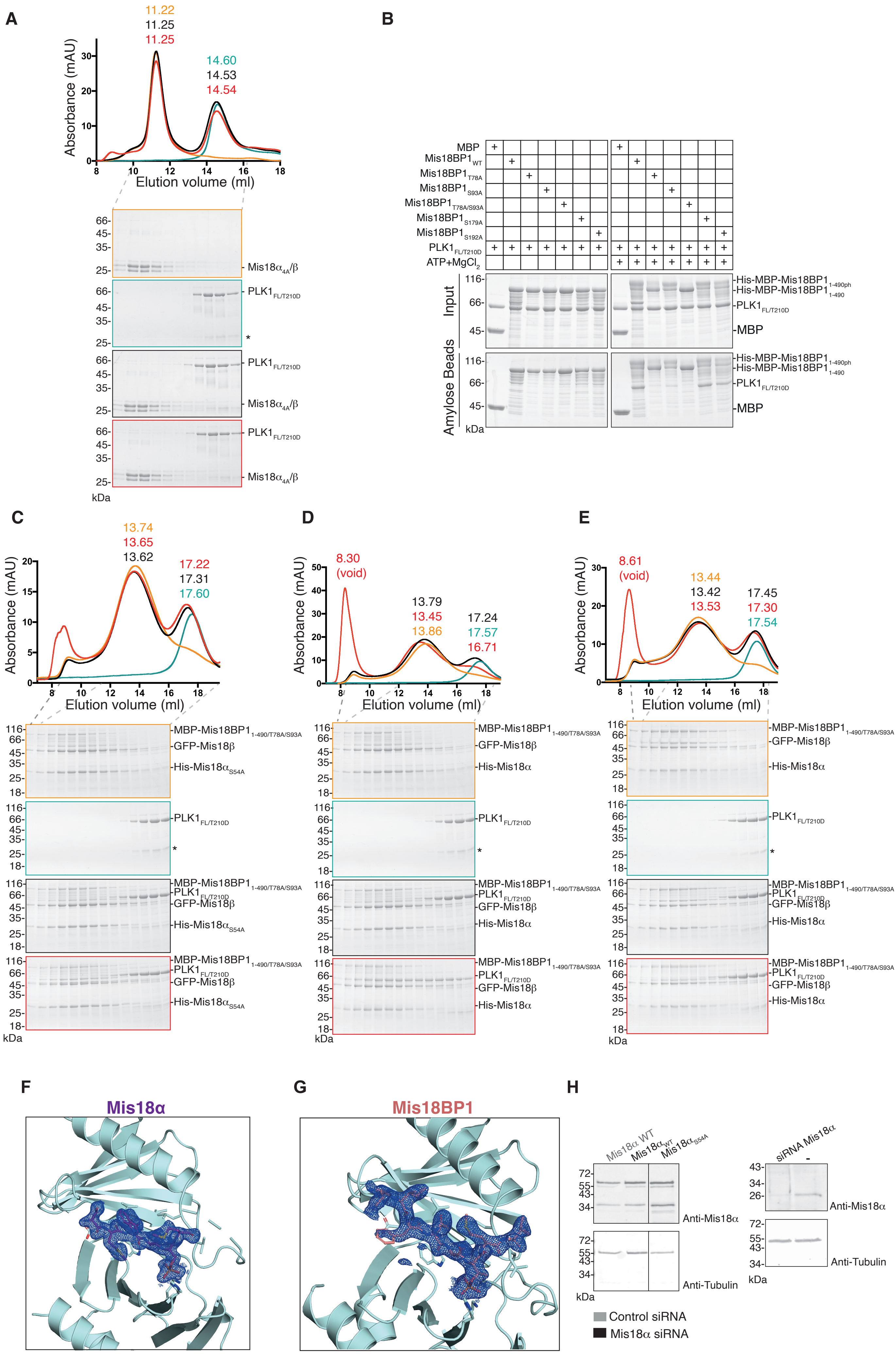

### Supplemental Figure 3

#### Supplementary 3

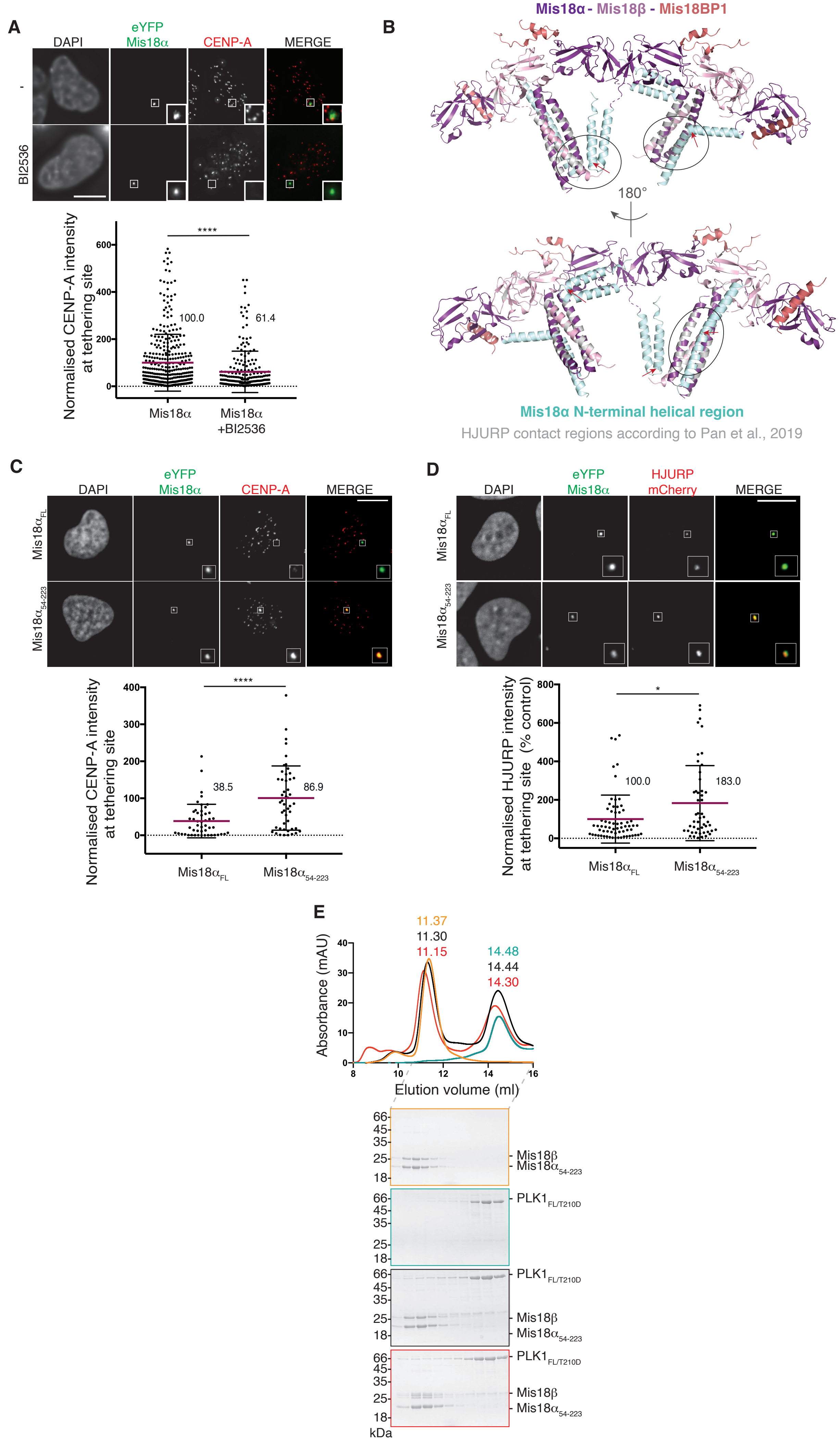

### Supplemental Figure 4

#### Supplementary 4

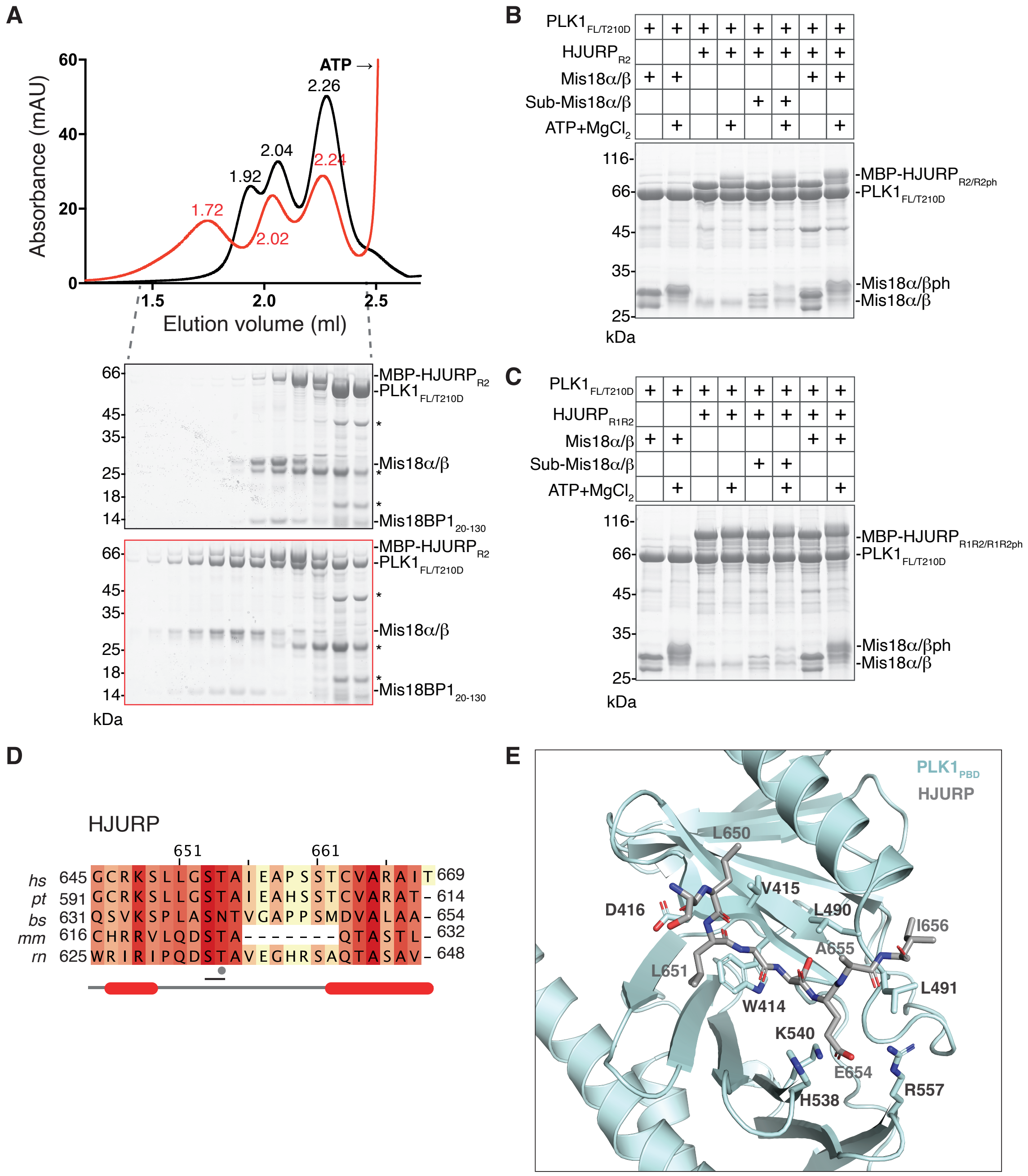
