## Supplemental Table 1 for "PLK1-Mediated Phosphorylation Cascade Activates the Mis18 Complex to Ensure Centromere Inheritance"

| Proteins | Positions within protein | PEP | Score | Phospho (ST) Probabilities |
|---|---|---|---|---|
| Mis18α_FL | 28 | 2.30E-09 | 162.49 | CS(0.015)DS(0.987)S(0.998)LLGKR |
| Mis18α_FL | 29 | ####### | 269.99 | CSDSS(1)LLGKR |
| Mis18α_FL | 36 | 1.05E-06 | 128.36 | RLS(0.969)EDS(0.011)S(0.02)R |
| Mis18α_FL | 39 | 3.41E-45 | 208.84 | RLSEDS(1)S(1)RHQLLQK |
| Mis18α_FL | 40 | ####### | 284.79 | RLSEDS(1)S(1)RHQLLQK |
| Mis18α_FL | 50 | 6.58E-04 | 63.522 | WAS(0.874)MWS(0.488)S(0.488)MS(0.15)EDASVADMER |
| Mis18α_FL | 53 | 6.58E-04 | 63.522 | WAS(0.837)MWS(0.779)S(0.288)MS(0.096)EDASVADMER |
| Mis18α_FL | 54 | 6.17E-04 | 59.252 | WAS(0.015)MWS(0.186)S(0.701)MS(0.087)EDAS(0.011)VADMER |
| Mis18α_FL | 56 | 6.35E-03 | 53.536 | WAS(0.065)MWS(0.624)S(0.479)MS(0.829)EDAS(0.003)VADMER |
| Mis18α_FL | 118 | 9.91E-29 | 168.83 | CVSCNVS(1)VDKEQK |
| Mis18α_FL | 187 | 1.45E-43 | 212.35 | QIVS(1)EDKELFNLESR |
| Mis18α_FL | 197 | 5.74E-12 | 135.55 | QIVSEDKELFNLES(1)R |
| Mis18α_FL | 204 | 7.23E-23 | 160.36 | VEIEKS(1)LTQMEDVLK |
| Mis18α_FL | 206 | 6.38E-04 | 99.941 | S(0.001)LT(0.999)QMEDVLK |
| Mis18α_FL | 227 | 5.55E-66 | 214.83 | LWEAESKLS(1)FATCKS |
| Mis18α_FL | 230 | 4.13E-05 | 125.23 | LSFAT(1)CKS |
| Mis18β_FL | 24 | 4.87E-04 | 108.72 | CATPPRGDFCGGT(1)ER |
| Mis18β_FL | 32 | 2.23E-16 | 127.74 | AIDQAS(0.999)FTTSMEWDTQVVK |
| Mis18β_FL | 34 | 7.56E-13 | 103.03 | AIDQAS(0.396)FT(0.591)T(0.012)S(0.001)MEWDTQVVK |
| Mis18β_FL | 35 | 3.03E-15 | 123.29 | AIDQAS(0.006)FT(0.039)T(0.782)S(0.173)MEWDTQVVK |
| Mis18β_FL | 36 | 8.92E-14 | 114.57 | AIDQAS(0.999)FT(0.001)T(0.076)S(0.881)MEWDT(0.043)QVVK |
| Mis18β_FL | 41 | 2.25E-13 | 115.89 | AIDQAS(0.884)FT(0.114)T(0.003)S(0.002)MEWDT(0.997)QVVK |
| Mis18β_FL | 183 | 8.05E-04 | 56.817 | AIVNAS(1)EMDIQNVPLSEK |
| Mis18β_FL | 221 | 2.93E-10 | 130.95 | ILS(0.005)EVT(0.995)PDQSKPEN |
| Mis18β_FL | 225 | 1.84E-36 | 179.9 | ILSEVTPDQS(1)KPEN |
| Mis18BP1 | 18 | 2.18E-06 | 103.97 | IYLPPEAS(0.983)S(0.017)QRR |
| Mis18BP1 | 19 | 6.24E-03 | 61.657 | IYLPPEAS(0.5)S(0.5)QRR |
| Mis18BP1 | 48 | 5.53E-03 | 53.034 | Y(0.709)QNS(0.146)S(0.146)LKLNDHKK |
| Mis18BP1 | 51 | 8.73E-07 | 194.06 | Y(0.003)QNS(0.998)S(0.998)LKLNDHKK |
| Mis18BP1 | 52 | ####### | 194.06 | YQNSS(1)LKLNDHK |
| Mis18BP1 | 67 | 5.35E-30 | 143.76 | NQFLKMT(1)T(1)FNNK |
| Mis18BP1 | 68 | 1.92E-03 | 76.17 | NQFLKMT(1)T(1)FNNK |
| Mis18BP1 | 81 | 3.31E-04 | 53.091 | NIFQS(0.164)T(0.202)MLT(0.249)EAT(0.076)T(0.081)S(0.099)N S(0.114)S(0.035)LDIS(0.979)AIKPNK |
| Mis18BP1 | 85 | 3.71E-04 | 54.514 | NIFQS(0.188)T(0.231)MLT(0.289)EAT(0.25)T(0.273)S(0.551)NS(0.182) S(0.055)LDIS(0.981)AIKPNK |
| Mis18BP1 | 86 | 1.88E-06 | 61.941 | NIFQS(0.188)T(0.231)MLT(0.289)EAT(0.25)T(0.273)S(0.551)NS(0.182) S(0.055)LDIS(0.981)AIKPNK |
| Mis18BP1 | 88 | 1.88E-06 | 61.941 | NIFQS(0.124)T(0.132)MLT(0.173)EAT(0.231)T(0.246)S(0.282)NS(0.41) S(0.41)LDIS(0.993)AIKPNK |
| Mis18BP1 | 89 | 1.88E-06 | 61.941 | NIFQS(0.124)T(0.132)MLT(0.173)EAT(0.231)T(0.246)S(0.282)NS(0.41) S(0.41)LDIS(0.993)AIKPNK |
| Mis18BP1 | 93 | 1.88E-06 | 61.941 | NIFQS(0.124)T(0.132)MLT(0.173)EAT(0.231)T(0.246)S(0.282)NS(0.41) S(0.41)LDIS(0.993)AIKPNK |
| Mis18BP1 | 135 | 3.06E-03 | 105.4 | NSS(1)LLEPQK |
| Mis18BP1 | 142 | 5.96E-03 | 74.376 | S(0.895)GNNET(0.102)FT(0.003)PNR |
| Mis18BP1 | 147 | 7.64E-08 | 122.46 | SGNNET(0.973)FT(0.027)PNR |
| Mis18BP1 | 191 | 4.85E-43 | 149.42 | ASVQGVPLES(0.761)S(0.239)NNDIFLPVK |
| Mis18BP1 | 192 | 7.32E-15 | 106.28 | ASVQGVPLES(0.212)S(0.788)NNDIFLPVK |
| Mis18BP1 | 240 | 3.59E-04 | 100.02 | NKT(1)LT(1)RAQLAK |
| Mis18BP1 | 242 | 3.59E-04 | 100.02 | NKT(1)LT(1)RAQLAK |
| Mis18BP1 | 253 | 1.24E-06 | 97.223 | QIFHS(1)KES(0.991)IVAT(0.007)T(0.002)K |
| Mis18BP1 | 256 | 7.48E-23 | 138.55 | QIFHSKES(1)IVATTK |
| Mis18BP1 | 260 | 7.48E-23 | 138.55 | QIFHS(0.999)KES(0.999)IVAT(0.991)T(0.011)K |
| Mis18BP1 | 261 | 5.44E-18 | 137.4 | QIFHS(0.093)KES(0.99)IVAT(0.961)T(0.957)K |
| Mis18BP1 | 263 | 1.69E-03 | 98.182 | ESIVAT(0.065)T(0.215)KS(0.72)K |
| Mis18BP1 | 299 | 7.66E-04 | 71.879 | NGS(1)LLMVSDSER |
| Mis18BP1 | 330 | 3.53E-08 | 79.283 | T(0.003)VPGET(0.996)GLPGS(0.001)MKDTCK |
| Mis18BP1 | 335 | 1.76E-02 | 41.825 | T(0.001)VPGET(0.033)GLPGS(0.966)MK |
| Mis18BP1 | 339 | 4.34E-05 | 70.783 | T(0.005)VPGET(0.946)GLPGS(0.204)MKDT(0.846)CK |
| Mis18BP1 | 352 | 2.18E-05 | 87.639 | LHIT(1)IPRR |
| Mis18BP1 | 362 | 3.05E-04 | 89.356 | RNIS(0.999)KLS(0.001)PPR |
| Mis18BP1 | 372 | 1.12E-48 | 162.93 | IFQT(1)VTNGLKK |
| Mis18BP1 | 374 | 1.97E-08 | 124.39 | IFQT(0.078)VT(0.922)NGLKK |
| Mis18α_FL | 26 | 3.22E-06 | 102.9 | GKCS(1)DS(0.259)S(0.741)LLGKR |
| Mis18α_FL | 28 | 3.14E-14 | 150.04 | CSDS(1)S(1)LLGKR |
| Mis18α_FL | 29 | 2.37E-19 | 171.63 | CSDS(1)S(1)LLGKR |
| Mis18α_FL | 36 | 2.75E-11 | 117.65 | RLS(1)EDS(1)S(1)RHQLLQK |
| Mis18α_FL | 39 | 4.70E-33 | 203.11 | RLS(1)EDS(1)S(1)RHQLLQK |

| | | | | |
|---|---|---|---|---|
| Mis18α_FL | 40 | 4.70E-33 | 203.11 | RLS(1)EDS(1)S(1)RHQLLQK |
| Mis18α_FL | 86 | 1.98E-20 | 124.8 | AQLEEEAAAAEERPLVFLCS(1)GCR |
| Mis18α_FL | 95 | 1.10E-33 | 165.48 | RPLGDS(1)LS(1)WVAS(0.006)QEDT(0.994)NCILLR |
| Mis18α_FL | 97 | 5.83E-09 | 79.9 | RPLGDS(1)LS(1)WVAS(0.006)QEDT(0.994)NCILLR |
| Mis18α_FL | 105 | 5.83E-09 | 79.9 | RPLGDS(1)LS(1)WVAS(0.006)QEDT(0.994)NCILLR |
| Mis18α_FL | 118 | 5.82E-42 | 214.47 | CVSCNVS(1)VDKEQK |
| Mis18α_FL | 126 | 8.52E-06 | 73.21 | CVSCNVS(0.002)VDKEQKLS(0.998)K |
| Mis18α_FL | 145 | 2.64E-06 | 76.573 | EKENGCVLET(0.009)LCCAGCS(0.991)LNLGYVYR |
| Mis18α_FL | 187 | 4.61E-06 | 74.519 | QIVS(1)EDKELFNLES(1)RVEIEK |
| Mis18α_FL | 197 | 1.96E-243 | 386.45 | QIVS(1)EDKELFNLES(1)RVEIEK |
| Mis18α_FL | 204 | 1.84E-100 | 280.54 | VEIEKS(1)LTQMEDVLK |
| Mis18α_FL | 206 | 1.47E-09 | 140 | SLT(1)QMEDVLK |
| Mis18α_FL | 224 | 2.89E-22 | 167.71 | LWEAES(1)KLS(0.999)FAT(0.001)CKS |
| Mis18α_FL | 227 | 3.61E-118 | 308.73 | LWEAESKLS(1)FATCK |
| Mis18α_FL | 230 | 5.40E-10 | 125.54 | LSFAT(0.998)CKS(0.002) |
| Mis18α_FL | 233 | 6.54E-05 | 52.784 | LWEAES(0.01)KLS(0.223)FAT(0.341)CKS(0.426) |
| Mis18β_FL | 10 | 2.27E-02 | 53.33 | S(0.999)RCAT(0.001)PPR |
| Mis18β_FL | 14 | 6.10E-04 | 55.44 | CAT(1)PPRGDFCGGTER |
| Mis18β_FL | 24 | 8.07E-10 | 99.02 | CATPPRGDFCGGT(1)ER |
| Mis18β_FL | 32 | 4.43E-52 | 212.34 | AIDQAS(1)FTT(0.013)S(0.987)MEWDTQVVK |
| Mis18β_FL | 34 | 4.90E-13 | 87.16 | AIDQAS(0.41)FT(0.59)T(0.523)S(0.477)MEWDT(0.001)QVVK |
| Mis18β_FL | 35 | 3.81E-33 | 178.88 | AIDQAS(1)FTT(0.699)S(0.3)MEWDTQVVK |
| Mis18β_FL | 36 | 4.43E-52 | 212.34 | AIDQAS(1)FTT(0.013)S(0.987)MEWDTQVVK |
| Mis18β_FL | 41 | 2.97E-18 | 130.56 | AIDQAS(0.939)FT(0.062)T(0.5)S(0.5)MEWDT(1)QVVK |
| Mis18β_FL | 68 | 3.12E-118 | 266.63 | GSSPLGPAGLGAEEPAAGPQLPS(1)WLQPER |
| Mis18β_FL | 100 | 5.19E-13 | 150.15 | S(1)LGAVVFSR |
| Mis18β_FL | 183 | 1.07E-77 | 254.02 | AIVNAS(1)EMDIQNVPLSEK |
| Mis18β_FL | 206 | 1.01E-03 | 74.46 | IVLT(1)HNRLKS(1)LMK |
| Mis18β_FL | 212 | 1.76E-03 | 49.34 | IVLT(1)HNRLKS(1)LMK |
| Mis18β_FL | 218 | 1.33E-03 | 52.56 | ILS(0.657)EVT(0.032)PDQS(0.311)KPEN |
| Mis18β_FL | 221 | 1.18E-117 | 300.71 | ILSEVT(1)PDQSKPEN |
| Mis18β_FL | 225 | 1.77E-117 | 299.25 | ILSEVTPDQS(1)KPEN |
| Mis18BP1 | 4 | 1.55E-215 | 361.83 | QTVDEALKDAQTNENLYFQSNAMIAT(0.906)PLKHS(0.094)R |
| Mis18BP1 | 9 | 2.74E-171 | 320.54 | DAQTNENLYFQSNAMIATPLKHS(1)R |
| Mis18BP1 | 18 | 1.72E-06 | 106.14 | IYLPPEAS(0.5)S(0.5)QRR |
| Mis18BP1 | 19 | 1.72E-06 | 106.14 | IYLPPEAS(0.5)S(0.5)QRR |
| Mis18BP1 | 33 | 3.31E-93 | 248.18 | NLPMDAIFFDS(1)IPSGTLTPVKDLVK |
| Mis18BP1 | 36 | 3.92E-10 | 248.18 | NLPMDAIFFDS(1)IPSGTLTPVKDLVK |
| Mis18BP1 | 51 | 0.000855468 | 71.379 | Y(0.001)QNS(0.999)S(1)LKLNDHKK |
| Mis18BP1 | 52 | 9.33E-67 | 241.3 | Y(0.001)QNS(0.999)S(1)LKLNDHKK |
| Mis18BP1 | 67 | 2.59E-51 | 235.73 | NQFLKMT(1)T(1)FNNK |
| Mis18BP1 | 68 | 2.26E-07 | 137.9 | NQFLKMT(1)T(1)FNNK |
| Mis18BP1 | 77 | 1.67E-06 | 66.381 | NIFQS(0.517)T(0.479)MLT(0.007)EAT(0.08)T(0.074)S(0.069)NS(0.06) S(0.056)LDIS(0.658)AIKPNKDGLK |
| Mis18BP1 | 78 | 9.22E-41 | 161.3 | NIFQS(0.168)T(0.831)MLT(0.001)EATTSNSSLDIS(1)AIKPNK |
| Mis18BP1 | 88 | 0.00011462 | 58.137 | NIFQS(0.085)T(0.091)MLT(0.114)EAT(0.144)T(0.156)S(0.169)NS(0.199) S(0.049)LDIS(0.992)AIKPNK |
| Mis18BP1 | 93 | 9.22E-41 | 161.3 | NIFQS(0.168)T(0.831)MLT(0.001)EATTSNSSLDIS(1)AIKPNK |
| Mis18BP1 | 131 | 8.03E-09 | 88.627 | DKQEQPS(1)RNS(1)S(1)LLEPQK |
| Mis18BP1 | 134 | 5.46E-14 | 111.57 | DKQEQPS(1)RNS(1)S(1)LLEPQK |
| Mis18BP1 | 135 | 5.46E-14 | 140.09 | DKQEQPS(1)RNS(1)S(1)LLEPQK |
| Mis18BP1 | 142 | 2.38E-08 | 89.541 | S(1)GNNET(0.998)FT(0.002)PNRVEK |
| Mis18BP1 | 147 | 4.59E-101 | 288.41 | SGNNET(1)FTPNR |
| Mis18BP1 | 161 | 1.37E-21 | 175.66 | KLQHT(1)YLCEEK |
| Mis18BP1 | 162 | 0.0003272 | 50.492 | LQHT(0.533)Y(0.555)LCEEKENNKS(0.9)FQS(0.01)DDS(0.001)S(0.001)LR |
| Mis18BP1 | 172 | 4.05E-40 | 159.59 | LQHT(0.949)Y(0.051)LCEEKENNKS(0.999)FQS(0.001)DDSSLR |
| Mis18BP1 | 191 | 5.63E-09 | 87.659 | ASVQGVPLES(0.855)S(0.145)NNDIFLPVK |
| Mis18BP1 | 192 | 5.63E-09 | 87.659 | ASVQGVPLES(0.5)S(0.5)NNDIFLPVK |
| Mis18BP1 | 218 | 5.20E-41 | 164.64 | APLHNLT(1)YELPTLNQEQENFLAVEAR |
| Mis18BP1 | 219 | 1.15E-09 | 82.476 | KAPLHNLT(0.685)Y(0.693)ELPT(0.622)LNQEQENFLAVEAR |
| Mis18BP1 | 223 | 5.20E-41 | 164.64 | APLHNLT(0.999)Y(0.001)ELPT(1)LNQEQENFLAVEAR |
| Mis18BP1 | 240 | 4.35E-10 | 142.1 | NKT(1)LT(1)RAQLAK |
| Mis18BP1 | 242 | 4.35E-10 | 142.1 | NKT(1)LT(1)RAQLAK |
| Mis18BP1 | 253 | 1.42E-11 | 123.72 | QIFHS(1)KES(1)IVAT(0.217)T(0.759)KS(0.024)K |
| Mis18BP1 | 256 | 1.39E-85 | 272.95 | QIFHS(1)KES(1)IVAT(0.217)T(0.759)KS(0.024)K |
| Mis18BP1 | 260 | 0.00154724 | 67.726 | ESIVAT(0.333)T(0.333)KS(0.333)K |
| Mis18BP1 | 261 | 1.35E-20 | 164.8 | QIFHS(0.001)KES(0.999)IVAT(0.013)T(0.987)K |
| Mis18BP1 | 263 | 3.25E-07 | 67.726 | S(0.494)KKDT(0.494)FVLES(0.022)VDS(0.045)ADEQFQNT(0.875) NAET(0.064)LS(0.005)T(0.002)NCIPIK |
| Mis18BP1 | 267 | 1.98E-31 | 133.07 | KDT(1)FVLESVDSADEQFQNTNAETLSTNCIPIK |
| Mis18BP1 | 272 | 2.64E-11 | 90.372 | S(0.333)KKDT(0.669)FVLES(0.565)VDS(0.422)ADEQFQNT(0.011) NAETLSTNCIPIK |

| | | | | |
|---|---|---|---|---|
| Mis18BP1 | 275 | 1.48E-07 | 74.806 | S(0.169)KKDT(0.169)FVLES(0.303)VDS(0.353)ADEQFQNT(0.006) NAETLSTNCIPIK |
| Mis18BP1 | 283 | 3.25E-07 | 63.277 | S(0.494)KKDT(0.494)FVLES(0.022)VDS(0.045)ADEQFQNT(0.875) NAET(0.064)LS(0.005)T(0.002)NCIPIK |
| Mis18BP1 | 299 | 7.19E-33 | 159.04 | NGS(1)LLMVSDSER |
| Mis18BP1 | 304 | 7.67E-14 | 101.73 | NGS(1)LLMVS(0.985)DS(0.561)ERT(0.343)T(0.084)EGT(0.022)S(0.006)QQK |
| Mis18BP1 | 306 | 7.19E-33 | 159.04 | NGS(1)LLMVS(0.006)DS(0.937)ERT(0.049)T(0.007)EGT(0.001)SQQK |
| Mis18BP1 | 309 | 2.97E-28 | 148.1 | NGS(1)LLMVS(0.057)DS(0.172)ERT(0.641)T(0.113)EGT(0.014)S(0.003)QQK |
| Mis18BP1 | 330 | 9.94E-17 | 116.98 | VPGET(1)GLPGSMKDTCK |
| Mis18BP1 | 335 | 3.22E-06 | 70.281 | TVPGET(1)GLPGS(0.793)MKDT(0.208)CK |
| Mis18BP1 | 339 | 1.60E-30 | 166.77 | TVPGETGLPGSMKDT(1)CK |
| Mis18BP1 | 352 | 0.000382086 | 99.973 | LHIT(1)IPRRS(1)K |
| Mis18BP1 | 357 | 0.0047981 | 53.034 | LHIT(1)IPRRS(1)K |
| Mis18BP1 | 362 | 3.58E-09 | 147.58 | NIS(1)KLSPPR |
| Mis18BP1 | 372 | 2.57E-51 | 228.62 | IFQT(1)VTNGLKK |
| Mis18BP1 | 374 | 5.50E-10 | 138.42 | IFQTVT(1)NGLKK |
| Mis18BP1 | 409 | 8.61E-41 | 189.42 | LIDVT(1)NIYWHSNVIIER |

### Supplementary Table 1. Phosphorylated Peptides
