## Supplemental Table 2 for "PLK1-Mediated Phosphorylation Cascade Activates the Mis18 Complex to Ensure Centromere Inheritance"

| | Mis18α | Mis18BP1 |
|---|---|---|
| **Data Collection** | | |
| Space group | P 1 21 1 | P 32 2 1 |
| **Cell dimensions** | | |
| *a*, *b*, *c*, (Å) | 43.1381, 56.5677, 49.5757 | 67.147, 67.147, 254.536 |
| α, β, γ (°) | 90, 115.738, 90 | 90, 90, 120 |
| Wavelength | 0.97625 | 0.91587 |
| Resolution (Å) | 44.66 – 1.94 (2.009-1.94) | 29.08 – 2.134 (2.211 – 2.134) |
| $R_{merge}$ | 0.1302 (0.8879) | 0.1043 (0.7229) |
| $R_{pim}$ | 0.08371 (0.7601) | 0.03468 (0.2468) |
| $I/\sigma I$ | 7.16 (0.50) | 15.79 (2.85) |
| Completeness (%) | 94.60 (61.63) | 99.16 (92.13) |
| Redundancy | 3.2 (1.9) | 10 (9.0) |
| **Refinement** | | |
| No. reflections | 48,573 (2172) | 377,150 (30,999) |
| $R_{work}$ (%) / $R_{free}$ (%) | 21.7 / 27.0 | 19.5 / 22.5 |
| **Average B** | | |
| Protein | 35.50 | 40.06 |
| **R.m.s deviations** | | |
| Bond length (Å) | 0.008 | 0.009 |
| Bond angles (°) | 1.10 | 1.27 |
| **Ramachandran values** | | |
| Favored (%) | 97.07 | 97.16 |
| Dissallowed (%) | 0 | 0.66 |

**Supplementary Table 2. Data Collection and Refinement Statistics.**
